# An RNA-binding region regulates CTCF clustering and chromatin looping

**DOI:** 10.1101/495432

**Authors:** Anders S. Hansen, Tsung-Han S. Hsieh, Claudia Cattoglio, Iryna Pustova, Xavier Darzacq, Robert Tjian

**Author notes:** ASH, TSH and CC contributed equally. Correspondence: XD and RT.

## Abstract

Mammalian genomes are folded into Topologically Associating Domains (TADs), consisting of cell-type specific chromatin loops anchored by CTCF and cohesin. Since CTCF and cohesin are expressed ubiquitously, how cell-type specific CTCF-mediated loops are formed poses a paradox. Here we show RNase-sensitive CTCF self-association *in vitro* and that an RNA-binding region (RBR) mediates CTCF clustering *in vivo*. Intriguingly, deleting the RBR abolishes or impairs almost half of all chromatin loops in mouse embryonic stem cells. Disrupted loop formation correlates with abrogated clustering and diminished chromatin binding of the RBR mutant CTCF protein, which in turn results in a failure to halt cohesin-mediated extrusion. Thus, CTCF loops fall into at least 2 classes: RBR-independent and RBR-dependent loops. We suggest that evidence for distinct classes of RBR-dependent loops may provide a mechanism for establishing cell-specific CTCF loops regulated by RNAs and other RBR partner.

## INTRODUCTION

Mammalian genomes are organized at multiple scales ranging from nucleosomes (hundreds of bp) to chromosome territories (hundreds of Mb) (Hansen et al., 2018a). At the intermediate scale of kilobases to megabases, mammalian interphase chromosomes are organized into local units known as Topologically Associating Domains (TADs) (Dixon et al., 2012; Nora et al., 2012). TADs are characterized by the feature that two loci within the same TAD contact each other more frequently, whereas two equidistant loci in adjacent TADs contact each other less frequently. Thus, TADs are thought to regulate contact probability between enhancers and promoters and therefore influence gene expression (Dekker and Mirny, 2016; Merkenschlager and Nora, 2016; Rowley and Corces, 2018).

Mechanistically, CCCTC-binding factor (CTCF) and the cohesin complex are hypothesized to form TADs through a loop extrusion mechanism: the cohesin ring complex entraps chromatin and extrudes intra-chromosomal chromatin loops until encountering convergently-oriented chromatin-bound CTCF molecules on both arms of the loop, halting cohesin-mediated extrusion (Alipour and Marko, 2012; Fudenberg et al., 2016, 2018; Ganji et al., 2018; Sanborn et al., 2015). CTCF and cohesin then hold together a TAD as a chromatin loop until these loop anchor proteins dissociate from chromatin. Thus, both loop extrusion and chromatin loop maintenance are dynamic processes (Fudenberg et al., 2016; Hansen et al., 2017, 2018a). Consistent with a causal role for CTCF and cohesin, TADs and chromatin loops largely disappear after acute depletion of CTCF and cohesin (Gassler et al., 2017; Nora et al., 2017; Rao et al., 2017; Schwarzer et al., 2017; Wutz et al., 2017). Moreover, CTCF and several cohesin subunits are among the most frequently mutated proteins in cancer (Hnisz et al., 2017; Lawrence et al., 2014), while disruption of TAD boundaries causes developmental defects (Lupianez et al., 2015).

The loop extrusion model can elegantly explain most experimental observations through a parsimonious mechanism (Fudenberg et al., 2018). In the model’s simplest form, any correctly oriented chromatin-bound CTCF should block cohesin-mediated loop extrusion. However, several conundrums remain. First, although almost all loops are anchored by CTCF, most CTCF-bound sites do not actually form loops (Merkenschlager and Nora, 2016; Rao et al., 2014). Why do only a subset of CTCF-sites form loops? Second, CTCF and cohesin are expressed in all cell types, but many TADs and loops display cell-type specific patterns and change during differentiation (Bonev et al., 2017; Pekowska et al., 2018; Stadhouders et al., 2018). How are new loops established and old ones disrupted during cell differentiation if CTCF and cohesin are always present? A potential solution to these unexplained features of TADs would be the existence of sub-classes of CTCF-boundaries, whose ability to block cohesin extrusion could be differentially regulated, though no such sub-classes have been described thus far.

Here, through an integrated approach combining genome editing, single-molecule and super-resolution imaging, *in vitro* biochemistry, ChIP-Seq and Micro-C, we identify critical functions of an RNA-binding region (RBR) in CTCF. Specifically, we show that the RBR mediates CTCF clustering and that loss of the RBR disrupts only a subset of CTCF-mediated chromatin loops. Our genome-wide analyses suggest that CTCF-boundaries can be classified into at least two sub-classes: RBR-dependent and RBR-independent. More generally, our work reveals a potential mechanism for establishing and maintaining specific CTCF loops, which may direct the establishment of cell-type specific chromatin topology during development.

## RESULTS

### CTCF self-associates in an RNA-dependent manner

We have previously shown that CTCF forms clusters in mouse embryonic stem cells (mESCs) and human U2OS cells (Hansen et al., 2017) and others have reported that CTCF forms larger foci in senescent cells (Zirkel et al., 2018). We therefore sought to investigate the molecular mechanisms underlying CTCF cluster formation. Clusters necessarily arise through direct or indirect self-association, so we took a biochemical approach to probe if and how CTCF self-interacts. Because CTCF over-expression causes artifacts and alters cell physiology (Hansen et al., 2017; Torrano et al., 2005), we used CRISPR/Cas9-mediated genome-editing to generate a mESC line in which one CTCF allele was 3xFLAG-Halo-tagged and the other allele V5-SNAPf-tagged (C62; Figure 1A-B). Consistent with CTCF clustering, when we immunoprecipitated (IP) V5-tagged CTCF, FLAG-tagged CTCF was pulled down along with it (coIP; Figure 1C; additional replicate and quantifications in Figure S1A-B). Conversely, immunoprecipitation of FLAG-tagged CTCF also co-precipitated significant amounts of V5-tagged CTCF (Figure S1C, Untreated). This observation using endogenously tagged CTCF confirms and extends earlier studies that observed CTCF self-association using exogenously expressed CTCF (Pant et al., 2004; Saldaña-Meyer et al., 2014; Yusufzai et al., 2004). But what is the mechanism of CTCF self-interaction? Benzonase treatment, which degrades both DNA and RNA (Figure S1D), strongly reduced the coIP efficiency (Figure 1C-D, S1A-C) whereas treatment with DNaseI had a significantly weaker effect on the CTCF self-coIP efficiency (Figure S1E). By contrast, treatment with RNase A alone severely impaired CTCF self-interaction (Figure 1C-D; S1A-C). We conclude that CTCF self-associates in a biochemically stable manner *in vitro* that is largely RNA-dependent while largely DNA-independent.

**Figure 1.**
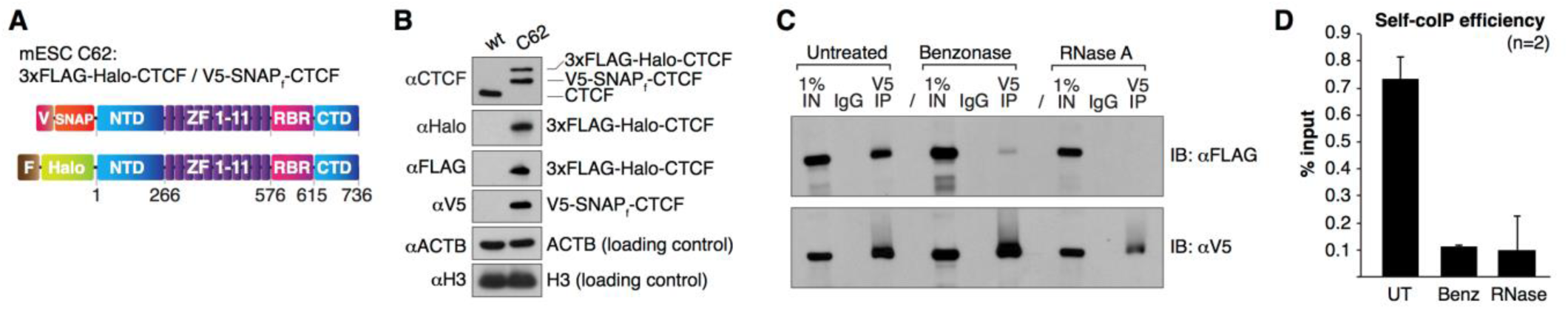
CTCF self-interacts in an RNA-dependent manner. (**A**) Overview of CTCF domains in the dual-tagged mESC clone C62. (**B**) Western blot of wild-type (wt) mESCs and endogenously dual-tagged C62 line showing that 3xFLAG-Halo-CTCF and V5-SNAP_f_-CTCF are similarly expressed and together roughly equal to CTCF levels in wt cells. (**C**) Representative coIP experiment indicating RNA-dependent CTCF self-interaction. Top: V5 IP followed by FLAG immunoblotting measures self-coIP efficiency (90% of the IP sample loaded); bottom: V5 IP followed by V5 immunoblotting controls for IP efficiency (10% of IP sample loaded). See also Figure S1A-E. (**D**) Quantification of CTCF self-coIP efficiency after normalization for V5 IP efficiency. Error bars are SD, n=2. See also Figure S1.

### An RNA-binding region (RBR) in CTCF mediates clustering

Our finding that CTCF self-association is predominantly RNA-mediated is perhaps surprising since CTCF is generally thought of as a DNA-binding protein. However, it confirms studies by Saldaña-Meyer *et al.*, who also showed that CTCF self-association depends on RNA but not DNA. Moreover, Kung *et al.* previously showed that CTCF binds RNA specifically and with high affinity (Kung et al., 2015). Importantly, Saldaña-Meyer *et al.* identified a short 38 amino acid RNA-Binding Region (RBR) C-terminal to CTCF’s 11-Zinc Finger DNA-binding domain that is necessary and sufficient for RNA binding and CTCF multimerization *in vitro* (Figure 1A)(Saldaña-Meyer et al., 2014). We therefore asked whether CTCF-clustering in cells is also RBR-dependent. The RBR largely corresponds to mouse CTCF exon 10, which we endogenously and homozygously replaced with a 3xHA tag in C59 Halo-CTCF mESCs (Hansen et al., 2017) to generate clone C59D2 ΔRBR (Halo-ΔRBR CTCF; Figure 2A-B and Figure S1F). ΔRBR-CTCF mESCs express a full length CTCF where most of the RBR (36 amino acids: N576 to D611) have been substituted with a short linker (GDGAGLINS) followed by a 3xHA tag, preserving the original exon 10 structure and length. Interestingly, while Halo-ΔRBR CTCF protein levels are only mildly reduced compared to Halo-wt-CTCF, as measured by TMR labeling and flow cytometry in live cells (Figure 1C; Figure S1G), ΔRBR-CTCF mESCs showed a ∼2-fold growth defect, suggesting that the RBR plays an important physiological role (Figure 2D). To test if the RBR mediates CTCF clustering, we performed super-resolution PALM imaging in fixed mESCs. We labeled Halo-CTCF with the PA-JF549 dye (Grimm et al., 2016), localized individual CTCF molecules inside the nucleus with a precision of ∼13 nm (Figure S1H) and reconstructed CTCF nuclear organization. Indeed, wt-CTCF (Figure 2E) showed noticeably higher clustering than ΔRBR-CTCF (Figure 2F), which we further verified and quantified using Ripley’s *L*-function (Besag, 1977; Boehning et al., 2018; Ripley, 1976) (Figure 2G; *L*(*r*)-*r* – values above 0 indicate clustering).

**Figure 2.**
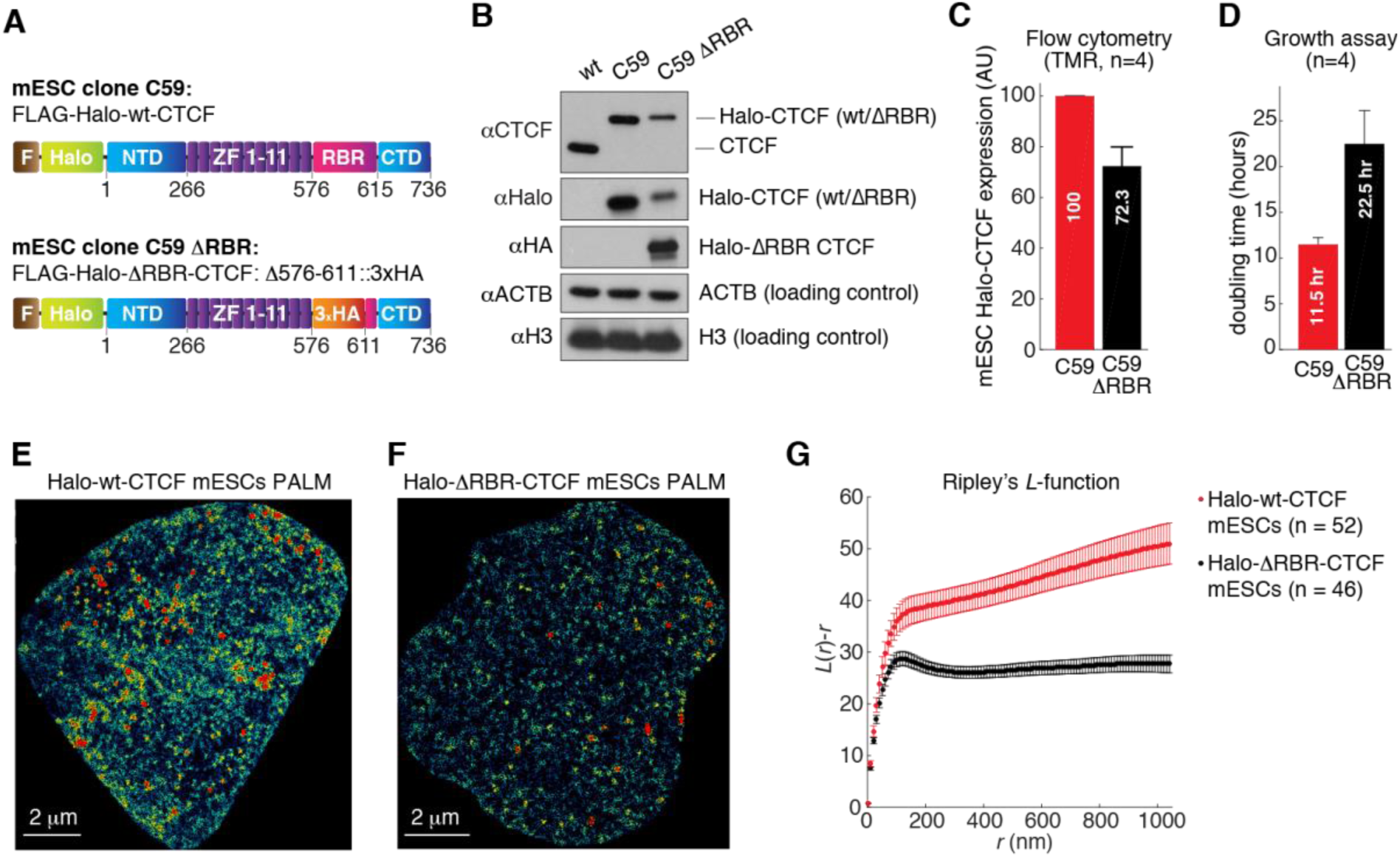
CTCF RBR region mediates CTCF clustering. (**A**) Overview of CTCF domains in the mESC clones C59 (Halo-wt CTCF) and C59 ΔRBR (Halo-ΔRBR CTCF). (**B**) Western blot of JM8.N4 wild type mESCs, C59, and C59 ΔRBR. (**C**) Flow cytometry measurement of Halo-CTCF abundance in live C59 Halo-wt CTCF and C59 ΔRBR mESCs after TMR labeling (mean and standard error). (**D**) Growth assay for C59 Halo-wt CTCF and C59 ΔRBR mESCs. Shows mean and standard error over 4 biological replicates. (**E-F**) Representative PALM reconstructions for Halo-wt CTCF (**E**) and Halo-ΔRBR CTCF (**F**). (**G**) Ripley’s L function for wt-CTCF (52 cells) and ΔRBR-CTCF mESCs (46 cells) (mean and standard error). See also Figure S1.

These results suggest that CTCF largely self-associates in an RNA-dependent manner and that CTCF clustering is significantly reduced, though not entirely abolished, in ΔRBR-CTCF mESCs. Although it is tempting to speculate that RNA(s) directly bind CTCF and hold together CTCF clusters *in vivo*, our results cannot distinguish between a mechanism where several CTCF proteins directly bind RNA from a model where CTCF indirectly interacts with an unknown factor, which then mediates CTCF-self-association in an RNase-sensitive manner.

### The CTCF RBR regulates 3D genome organization, but not compartments

In a companion paper focused on the biophysics of nuclear sub-diffusion (Hansen et al., 2018b), we show using single-particle tracking experiments that CTCF exhibits unusual anisotropic diffusion: once CTCF has moved in one direction, it is substantially more likely to move backwards, than to continue forwards. Our theoretical work suggests that this is due to transient trapping in localized zones/domains of a characteristic size (∼200 nm). Specifically, this local trapping is largely lost for ΔRBR-CTCF, suggesting that the local zones/domains may correspond to CTCF clusters (Figure 2D-F). We term this mechanism Anisotropic Diffusion through transient Trapping in Zones (ADTZ). Moreover, we show that CTCF’s cognate DNA-target search mechanism is RBR-guided, such that ΔRBR-CTCF takes ∼2.5-fold longer to locate a cognate DNA-binding site. We provide full details on the theory and analysis of the diffusion mechanism in our companion paper (Hansen et al., 2018b). Here we focus on the primary function of CTCF, which is to regulate 3D genome organization. We therefore next investigated whether impaired CTCF clustering, self-association, and target searching of ΔRBR-CTCF (Figures 1-2; (Hansen et al., 2018b)) might also impact 3D genome organization, using a high resolution genome-wide chromosomal conformation capture (3C) assay, Micro-C. Unlike Hi-C which uses restriction enzymes, Micro-C fragments chromatin to single nucleosomes using micrococcal nuclease and generates 3D contact maps of the genome at all biologically-relevant resolutions (Hsieh et al., 2016, 2015). Originally developed for analyzing the small yeast genome, here we have adapted a Micro-C protocol for large-genome organisms and successfully recapitulated all the chromatin features previously identified by Hi-C (unpublished manuscript by T.S.H *et al.*). We applied this Micro-C protocol to C59 (wt-CTCF) and C59D2 (ΔRBR-CTCF) mESCs (Figure 2A) over two replicates and generated ∼668 and ∼694 million unique contacts, respectively. Moreover, we also performed CTCF and Cohesin (Smc1a) chromatin immunoprecipitation followed by DNA sequencing (ChIP-Seq) in two replicates for wt-CTCF and ΔRBR-CTCF mESCs (see below). We then surveyed 3D genome organization and analyzed features across several scales (Figure 3A) including compartments, topologically-associating domains (TADs), loops, and stripes (Fudenberg et al., 2018). We readily detected clear changes in the genome organization of ΔRBR-CTCF mESCs relative to wt-CTCF at multiple chromosomal scales (Figure 3A), and began our analysis at the large end of the scale: compartments.

**Figure 3.**
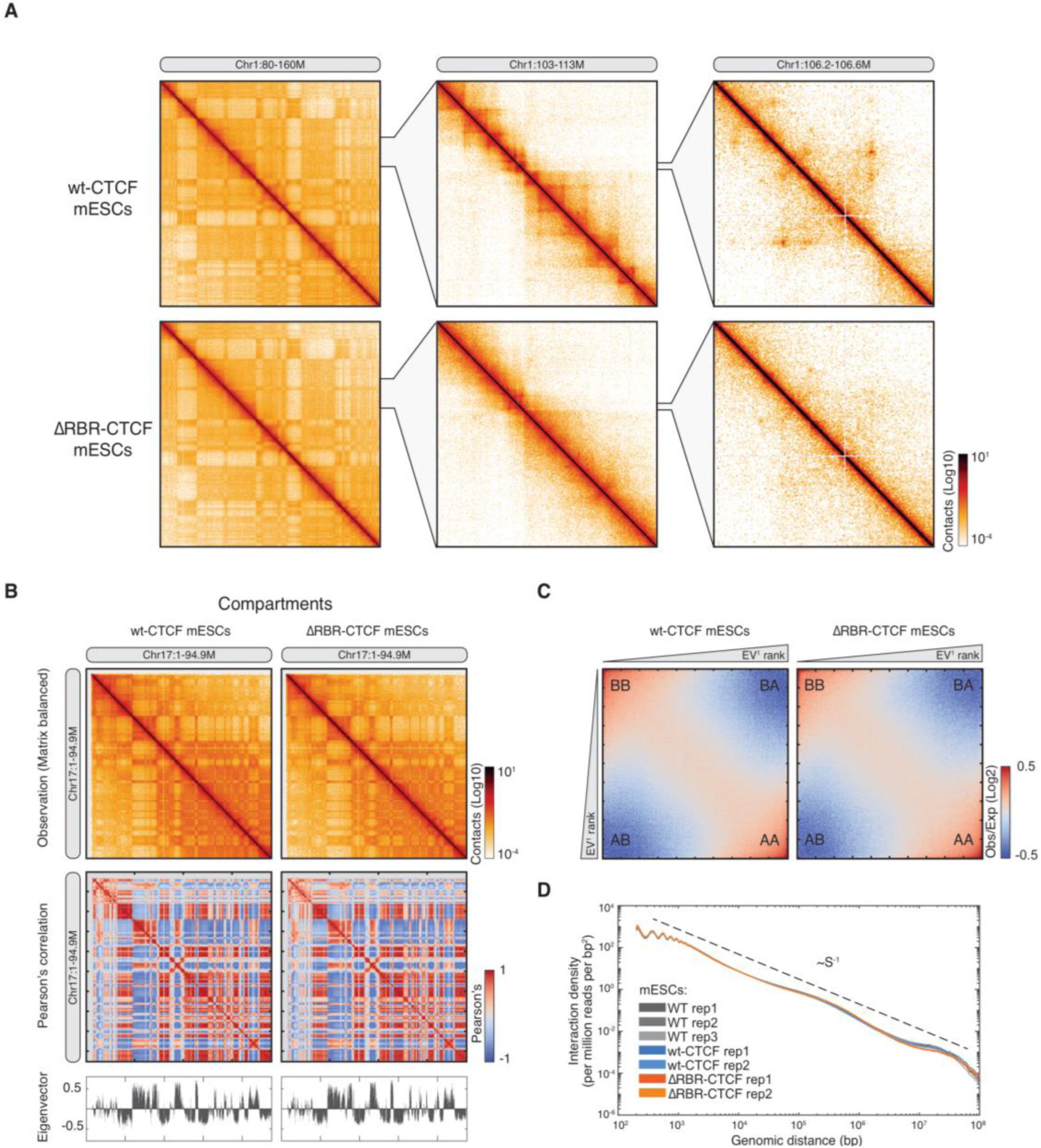
Compartments are largely unchanged in ΔRBR-CTCF mESCs. (**A**) Overview of Micro-C contact matrices at multiple resolutions in wt-CTCF and ΔRBR-CTCF mESCs. Contact matrix normalization: iterative correction and eigenvector decomposition (ICE); color scale: log10 unless otherwise mentioned. The differences between wt-CTCF and ΔRBR-CTCF mESCs can be observed at sub-compartmental levels as shown in the middle and right panels. (**B**) Comparison of chromosome compartments. An example of plaid-like chromosome compartments at Chr17 is shown as ICE balanced contact matrices, Pearson’s correlation matrices, and Eigenvector analysis for the first principle component at 100-kb resolution. There is no significant alteration in global active and inactive chromatin compartments between wt-CTCF and ΔRBR-CTCF mESCs. (**C**) Saddle plot for compartmentalization strength. The plot was calculated by the average distance-normalized contact frequencies between 100-kb bins *in cis* with ascending eigenvector values (EV^1^). All distance-normalized contact matrices in this study are shown at log2 scale between the scale at −1 to 1 or −0.5 to 0.5 unless otherwise mentioned. In the saddle plot, the upper-left and bottom-right represent the contact frequency between B-B and A-A compartments and upper-right and bottom-left show the frequency of inter-compartment interactions. (**D**) Genome-wide contact probability scaling plot. The scaling curves are plotted by the interaction density (per million reads per bp^2^) against genomic distance from 100bp to 100Mb. Curves for biological replicates of wt-CTCF and ΔRBR-CTCF mESCs overlap and decay at slope of −1, as previously reported (Lieberman-aiden et al., 2009). Due to potential artifacts introduced by fragment self-ligation, we did not consider reads below 100 bp.

Mammalian chromosomes can be divided into two major compartments (Lieberman-aiden et al., 2009): A-compartments, composed mainly of active euchromatin and B-compartments, composed mainly of inactive and gene-poor heterochromatin and lamina-associated domains (van Steensel and Belmont, 2017). We observed no significant change in compartmentalization when comparing wt-CTCF and ΔRBR-CTCF mESCs (Figure 3B), nor did we observe significant changes in A-A, A-B, B-A or B-B contact frequency (Figure 3C). Moreover, averaged over the whole genome, we observed the same contact-probability scaling with genomic distance for wt-CTCF and ΔRBR-CTCF mESCs (Figure 3D). We conclude that the CTCF RBR does not affect the global distribution of active and inactive chromatin, and note that this result is consistent with previous studies that also recorded no strong changes in compartmentalization even after near-complete CTCF degradation (Nora et al., 2017; Wutz et al., 2017).

### Loss of CTCF RBR disrupts a subset of TADs

Having analyzed compartments, we next zoomed in and analyzed TADs. TADs are demarcated by a pair of strong boundaries, or insulators, which are frequently bound by the architectural proteins CTCF and cohesin, and typically span lengths of ∼100 kb to ∼1 Mb in mouse and human genomes (Merkenschlager and Nora, 2016; Rowley and Corces, 2018). TADs are characterized by the feature that two loci inside the same TAD contact each other more frequently than two equidistant loci in different TADs (Dixon et al., 2012; Nora et al., 2012). We defined TADs using either arrowhead or insulation score (Crane et al., 2015; Rao et al., 2014) and arbitrarily chose a cut-off value to obtain ∼3500 TADs in wt-CTCF mESCs, corresponding to the previously reported TAD size and number (Forcato et al., 2017). Although the inferred number and size of TADs depends on the algorithm and the resolution of the maps (Forcato et al., 2017), we generally observed fewer and larger TADs in ΔRBR-CTCF mESCs (Figure 4A and Figure S2A). In brief, our insulation analysis called 3,666 and 2,793 TADs with average TAD sizes of ∼715 kb and ∼936 kb in wt-CTCF and ΔRBR-CTCF mESCs, respectively (Figure 4B). To gain further insight into how TAD organization may be altered in ΔRBR-CT CF mESCs, we resized and aggregated over all TADs genome-wide (Figure 4C). TADs were clearly weaker in ΔRBR-CTCF mESCs and also showed substantially weaker insulation strength (Figure 4D and Figure S2B-C), which was reduced by ∼19% at the center of wild-type TAD boundaries. Interestingly, however, we also observed 604 new insulators in ΔRBR-CTCF mESCs, that we will refer to as “cryptic insulators” (Figure 4B). To further analyze these cryptic insulators, we calculated their relative distance to the original ones (Figure S2D) and found that they are usually shifted by ∼200kb from the original insulator. A portion of cryptic insulators correspond to new cohesin (Smc1a) peaks that appeared either upstream or downstream of the disrupted CTCF/cohesin binding sites (Figure 4E, pink arrow in browser track b). We analyze these cryptic insulators and cohesin peaks in greater detail below.

**Figure 4.**
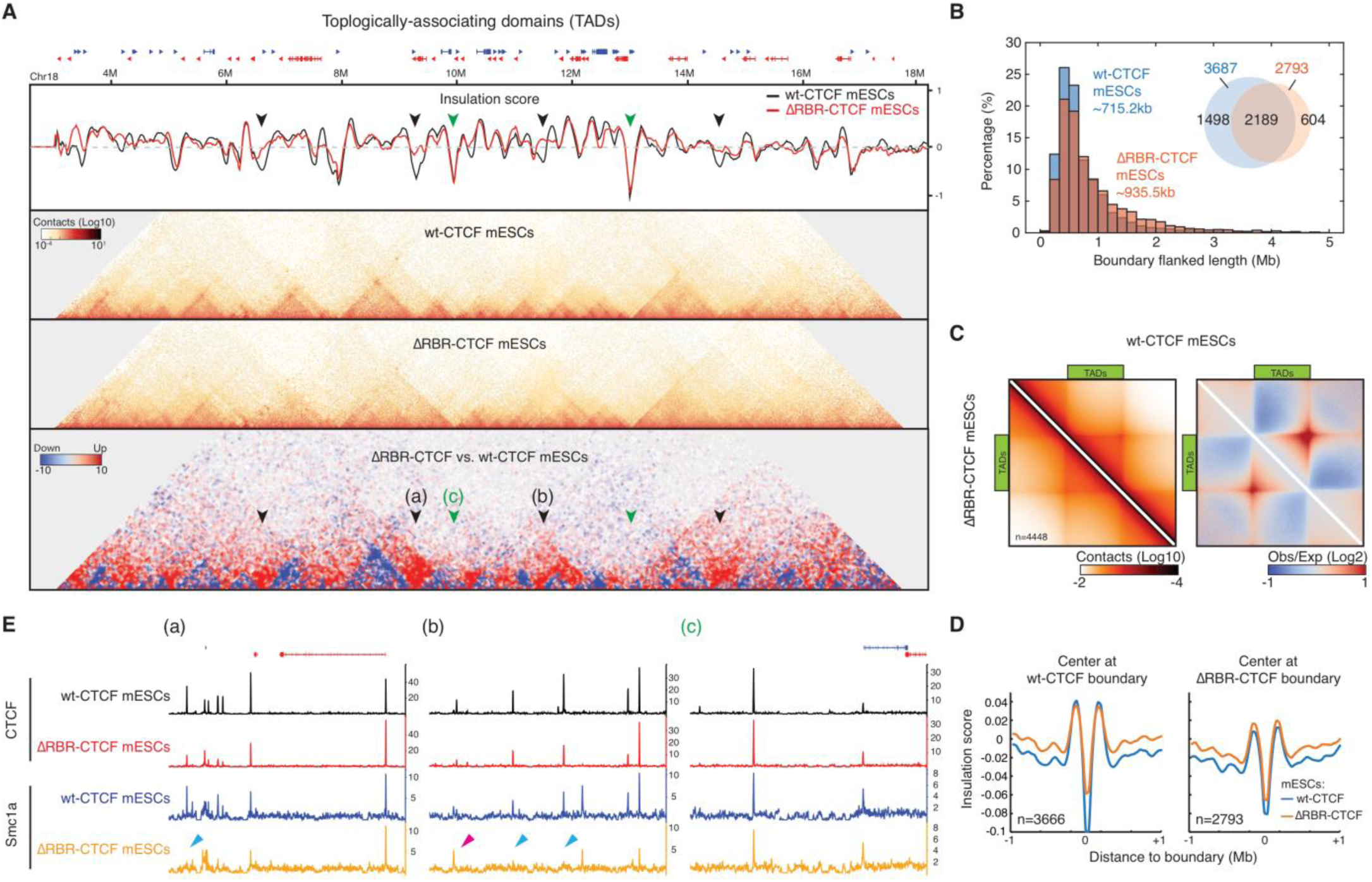
TAD organization is significantly changed in ΔRBR-CTCF mESCs. (A) An example of TAD/boundary disruption in ΔRBR-CTCF mESCs. A snapshot of insulation score curves, 45°-rotated contact maps, and differential contact matrix (from top to bottom) were plotted for Chr18: 3M-18M. The insulation score was calculated by a 200-kb sliding window in 20-kb resolution maps. A lower value of insulation score means stronger insulation strength. The black arrows indicate examples of loss of insulation in ΔRBR-CTCF mESCs and the green arrows indicate the unaffected insulators. The differential contact matrices were generated by subtraction of the normalized ΔRBR-CTCF matrix to wt-CTCF matrix. Blue triangles along the diagonal indicate weaker TADs in ΔRBR-CTCF mESCs. Red indicates “bleed-through”, i.e. loss of TAD insulation (black arrows). (**B**) Size distribution of TADs. The histogram shows the size distribution of boundary/insulator-flanked regions in wt-CTCF and ΔRBR-CTCF mESCs. ΔRBR-CTCF mESCs lose ∼15% of TADs at size smaller than 100kb but exhibit more TADs at size over 150kb scale. Inset: Venn diagram. ΔRBR-CTCF mESCs lose 1,474 out of 3,666 insulators identified in wt-CTCF mESCs but gain 604 cryptic insulators. (**C**) Aggregate peak analysis for TADs. TADs in wt-CTCF mESCs were identified through an additional TAD calling algorithm (arrowhead) and rescaled and aggregated (n=4,448) at the center of plot with ICE normalization (Left) or distance normalization (Right). The wt-CTCF is shown on the top half and ΔRBR-CTCF is shown on the bottom half. (**D**) Genome-wide averaged insulation plotted vs. distance around insulation center. Insulation strength is weaker in ΔRBR-CTCF mESCs when centering at wild-type insulators, but there is no significant change when centering at ΔRBR-CTCF insulators. (**E**) Browser tracks. ∼200kb snapshot regions around the arrows (a, b, and c) indicated in (**A**) are shown with CTCF and cohesin (Smc1a) ChIP-Seq data. Panel (a) and (b) display regions with strong depletion of insulation in ΔRBR-CTCF mESCs and panel (c) shows a region with little effect. The blue arrows indicate examples for loss of Smc1a peaks in ΔRBR-CTCF mESCs and the pink arrow indicates an example for gain/shift of Smc1a peak. **See also Figure S2 and S3.**

We next inspected local regions that were altered in ΔRBR-CTCF mESCs, superimposing Micro-C and ChIP-Seq results. Of note, when using spike-in normalization for ChIP-Seq analysis, ΔRBR-CTCF signal appeared globally reduced compared to wt-CTCF, while Smc1a binding was largely unaltered at preserved sites (∼60% of wt Smc1a binding sites; Figure S5B-C, Figure 6D). Because biochemical experiments showed reduced stability of the ΔRBR-CTCF protein after cell lysis (Figure 6A), we could not determine whether the dampened ChIP-Seq signal resulted from reduced ChIP efficiency, diminished genomic occupancy of ΔRBR-CTCF, or both. We thus decided to normalize data by sequencing depth instead, and avoid direct comparisons between wt-CTCF and ΔRBR-CTCF ChIP-Seq signals to draw conclusions. When inspecting local genomic regions, we noticed that CTCF and cohesin (Smc1a) binding was strongly depleted at some specific loci in the ΔRBR-affected boundary (Figure 4E, blue arrows in browser track (a) and (b)). Conversely, CTCF and cohesin binding was largely retained at unaffected boundaries (Figure 4E, browser track c). We conclude that the RBR contributes significantly to CTCF’s role in forming TADs. This is unlikely an indirect effect, because: i) the cell cycle phase distribution was identical between wt-CTCF and ΔRBR-CTCF mESCs, despite the growth defect of the latter (Figure S3A-B) and, ii) although the ΔRBR-CTCF expression level was somewhat lower (reduced by 28%) compared to wt-CTCF (Figure 1C and Figure S1G), Nora *et al.* previously demonstrated that TAD organization in mESCs is preserved for the most part even after 85% reduction of CTCF levels (Nora et al., 2017).

**Figure 6.**
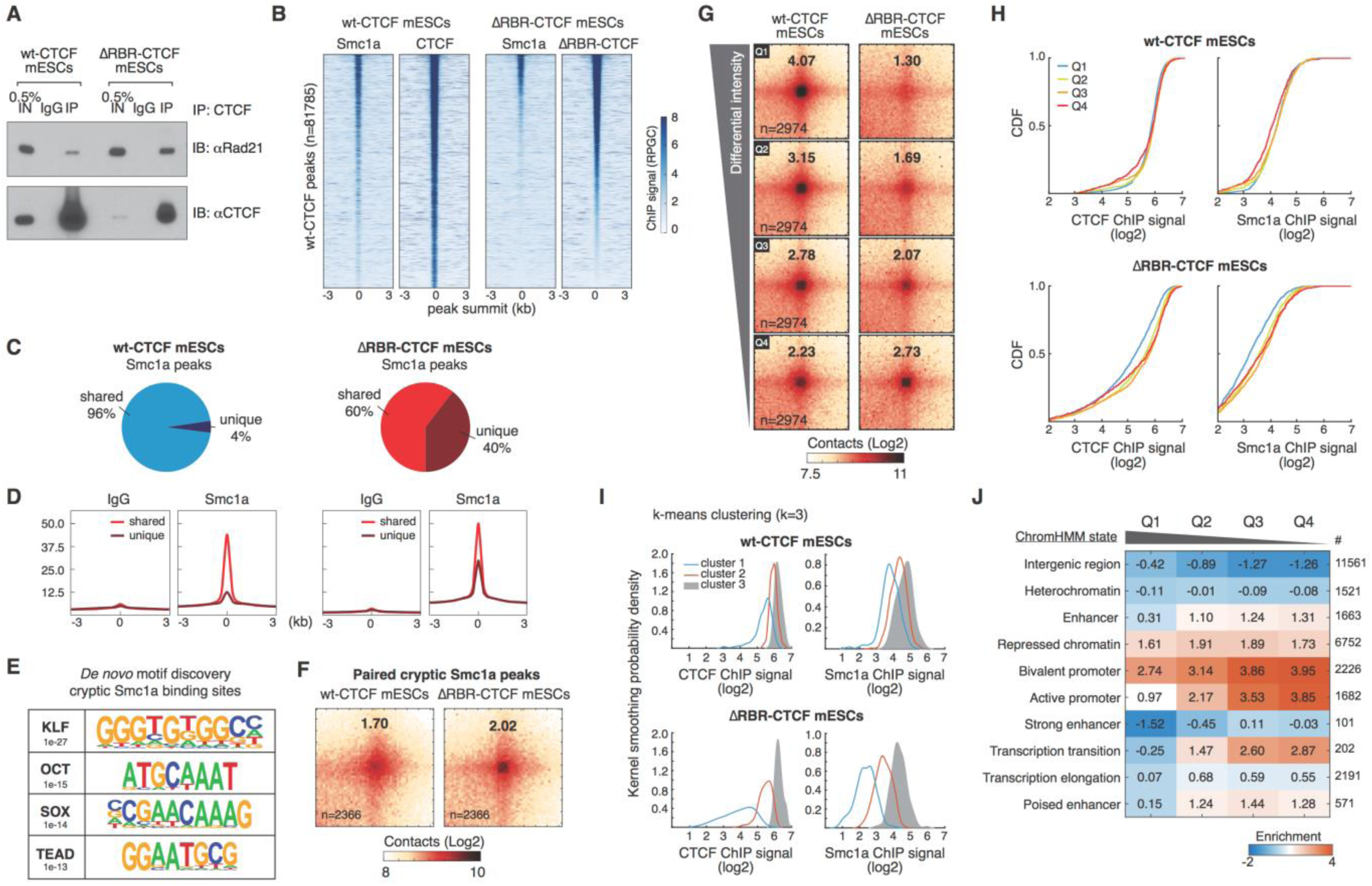
ΔRBR-CTCF still interacts with cohesin and loops lost in ΔRBR-CTCF mESCs have less CTCF and cohesin bound. (**A**) Representative coIP experiment showing that ΔRBR-CTCF stills interacts with cohesin. CTCF antibodies can pull down Rad21 cohesin subunit in both wt- and ΔRBR-CTCF mESCs. (**B**) Heatmaps of CTCF and Smc1a ChIP-Seq signal (deepTools RPGC: reads per genomic content) around wt-CTCF peaks as called by MACS2. Heatmaps are sorted by ΔRBR-CTCF peak intensity. (**C**) Fraction of unique and shared Smc1a peaks in wt (left, blue) and ΔRBR (right, red) mESCs, as called by MACS2. Shared peaks are those overlapping at least 1 bp with a CTCF peak in wt-mESCs and/or an Smc1a peak in the other cell line. (**D**) Mean ChIP-Seq signal intensity (scaled by number of reads after spike-in normalization; see STAR methods) of IgG and Smc1a ChIP-Seq signal in wt (left) and ΔRBR (right) mESCs around Smc1a peaks called by MACS2 in ΔRBR mESCs as shared or unique as described in (**C**). (**E**) *De Novo* motif discovery around cryptic Smc1a peaks. Searching protocol: identify significant motifs in size 8 – 13 bp within ±250 bp of the cryptic peaks. (**F**) Aggregate peak analysis for the cryptic Smc1a peaks. All the cryptic Smc1a peaks were paired (n=17,594 peaks) and pairs further than 1Mb excluded. Loops were selected at 5kb resolution using FDR < 0.1 (n=2366 pairs). (**G**) Aggregate peak analysis for differential loop intensity. Loops were sorted into four quartiles based on differential loop intensities between wt-CTCF and ΔRBR-CTCF mESCs. 2,974 loops in each quartile were aggregated at the center of a 50kb window and quantified as above. (**H**) Cumulative distribution function (CDF) curves of ChIP enrichment at the loop anchors. Loop anchors were identified as described in methods. CTCF and Smc1a ChIP signals were quantified as the log2 enrichment around ±250 bp of the anchor center. (**I**) *k*-means clustering analysis of CTCF and cohesin (Smc1a) ChIP-Seq data in the Q1 loop anchors. The filtered Q1 loop anchoring sites were further analyzed by *k*-means clustering (*k*=3). The outputs of clustering analysis were plotted as kernel smoothed histograms. Heatmaps with the peaks at the center across a 3kb region are shown in Figure S5F. (**J**) Enrichments of genomic features at loop anchors. ChromHMM analysis for mouse mm10 genome was from (Bogu et al., 2016). The heatmap is shown as log2 enrichment of the loop anchors in each chromatin state. Note that Q1 loops are largely depleted in most chromatin states and only slightly enriched in H3K27me3 chromatin. See also Figure S5 and S6.

### CTCF loops fall into RBR-dependent and independent subclasses and loss of the CTCF RBR causes longer stripes

Many TADs show corner peaks of Micro-C signals at their summit, suggesting that they are held together as loop structures (Fudenberg et al., 2018; Rao et al., 2014) (see also Figures 3A and 4C). According to the loop extrusion model (Fudenberg et al., 2016, 2018; Sanborn et al., 2015), cohesin extrudes DNA loops until it is stopped by pairs of chromatin-bound CTCF proteins in a convergent orientation. However, it is unclear which protein domain(s) in CTCF are required to block cohesin extrusion. To test whether the RBR plays any role in loop formation and/or maintenance, we analyzed the contact maps at high resolution (∼1 kb to 5 kb) and identified ∼14,372 loops in wt-CTCF mESCs by the method described by Rao *et al.* (Rao et al., 2014). Overall, out of 14,372 called loops, 57% (8,189 loops) were weakened by at least 1.5-fold in ΔRBR-CTCF mESCs and 39% (5,490) by at least 2-fold relative to wild type (Figure 5A-B). We next performed genome-wide loop aggregation analysis. The loop strength in C59 wt-Halo-CTCF mESCs is about as strong as in mESCs with untagged CTCF (Bonev et al., 2017) (Figure S4A), confirming that our endogenously tagged Halo-CTCF mESCs behave as wild-type mESCs (Hansen et al., 2017). However, the loop strength was greatly reduced in ΔRBR-CTCF mESCs (Figure 5C and Figure S4A). As a comparison, we re-analyzed Hi-C data at loops in mESCs with a CTCF degron from Nora *et al.* (Nora et al., 2017) and found that the loss in loop strength upon near-complete CTCF degradation is actually comparable to the defect in loop strength we observe for ΔRBR-CTCF mESCs (Figure S4A-C). Although technical differences between Micro-C and Hi-C make a direct comparison difficult, these results nevertheless underscore how severe the loop strength defect is in ΔRBR-CTCF mESCs.

**Figure 5.**
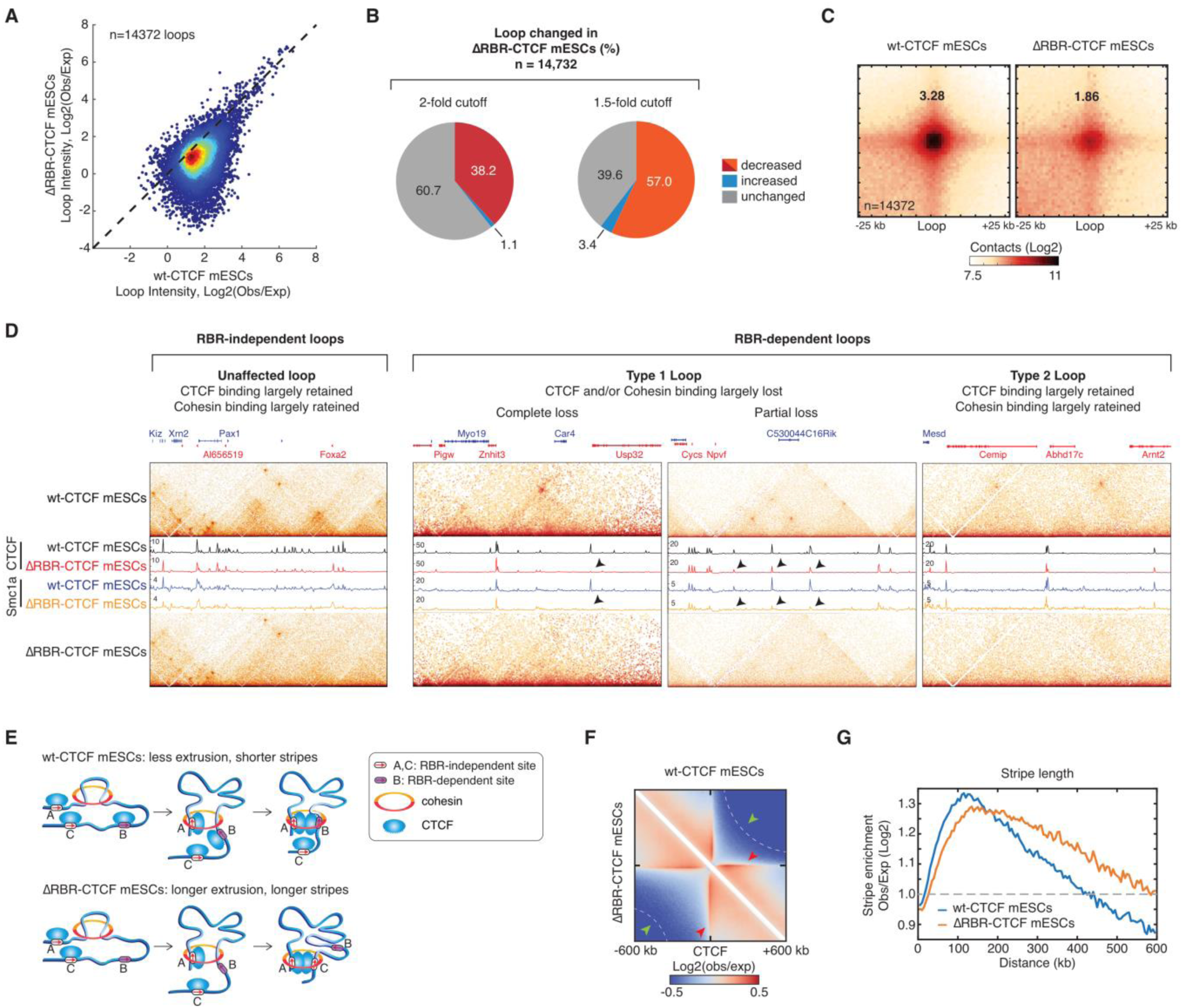
Genome organization at the level of both loops and stripes is altered in ΔRBR-CTCF mESCs. **(A)** Scatter plot showing individual loop intensities in wt-CTCF vs. ΔRBR-CTCF mESCs. 14,372 loops were identified in wt-CTCF mESCs with a false discovery rate < 0.1. Loop intensity was calculated as log2 enrichment of center pixel over expected bottom-left pixels in 1 kb, 5 kb, or 10 kb resolution. (**B**) Pie charts showing affected loops. Approximate 8,189 loops are decreased by at least 1.5-fold and 5,490 loops are decreased by at least 2-fold in ΔRBR-CTCF compared to wt-CTCF mESCs. Some loops are not affected or even strengthened in ΔRBR-CTCF mESCs. (**C**) Aggregate peak analysis for loops. The called loops were aggregated at the center of a 50-kb window in 1kb resolution. The genome-wide averaged loop enrichment was calculated by the fold enrichment (center pixel/expected bottom-left pixels). (**D**) Snapshots of four genomic regions representative of different CTCF loop types. Zoomed-in contact maps were plotted on the top and bottom panels for wt-CTCF and ΔRBR-CTCF mESCs, respectively. CTCF and cohesin (Smc1a) ChIP-Seq data is overlaid. From left to right, shown are examples of RBR-independent loops and of the two subtypes of RBR-dependent loops (with two examples of partial and complete loss of CTCF/cohesin binding for loop type 1). (**E**) Loop extrusion sketch. Illustration of why loss of a subset of CTCF boundaries would result in longer stripes within the context of the loop extrusion model (Fudenberg et al., 2018). (**F**) Aggregation plot centered at top CTCF peaks. The contact matrices were aggregated around the top 10,000 CTCF ChIP-Seq peaks using a ±600kb window. wt-CTCF mESCs are shown on the top half of plot and ΔRBR-CTCF mESCs are shown on the bottom half. Red arrows indicate stripes/flames. Green arrows and white dash lines indicate insulation strength that represents the capacity to prevent interactions across the boundary. (**G**) A quantitative curve for stripe length. Stripes enrichments were calculated in log2 ratio of observed over expected contacts. The value of significant enrichment was preset as a threshold with 2-fold enrichment labeled in gray dash line on the plot. CTCF-mediated stripes are extended by ∼200 kb in ΔRBR-CTCF mESCs. See also Figure S4.

Surprisingly, the effect of deleting the RBR was highly heterogeneous: some CTCF loops were unaffected or even strengthened, while others were significantly weakened or completely disrupted in ΔRBR-CTCF mESCs (Figure 5D). Qualitatively, we could distinguish two general categories of loops: an RBR-independent class (Figure 5D, left) and an RBR-dependent class (Figure 5D, right). When we overlaid the ChIP-Seq tracks on the Micro-C contact maps, we noticed that CTCF and cohesin (Smc1a) binding was largely preserved at the anchors of RBR-independent loops, as expected. However, we could distinguish at least two sub-types of loops which were lost in ΔRBR-CTCF mESCs: 1) partial or complete loss of ΔRBR-CTCF and/or cohesin binding at least at one loop anchor (Figure 5D, type 1 loops); 2) no significant change in either ΔRBR-CTCF or cohesin binding (Figure 5D type 2 loops). Thus, whereas loop loss for type 1 loops can be explained through loss of CTCF and cohesin binding, differential changes in CTCF and cohesin binding cannot readily explain loss of type 2 loops. We discuss the mechanistic implications of these findings in greater detail below.

Finally, we analyzed stripes/flames (Fudenberg et al., 2018). Stripes are thought to be a signature for loop extrusion and are hypothesized to occur when cohesin engaged in extrusion reaches one CTCF boundary before the other (Figure 5E). In particular, since we see loss of a subset of CTCF loops and ∼200 kb larger TADs in ΔRBR-CTCF mESCs (Figure 4B and S2D), a prediction of the loop extrusion model would be that an extruding cohesin must on average travel longer to reach a functional CTCF site in ΔRBR-CTCF mESCs, and therefore, we should see longer stripes (Figure 5E). To test this prediction, we compiled contact matrices using the top 10,000 wt-CTCF ChIP signals at the center of the plot and found that the stripes in ΔRBR-CTCF mESCs are less intense at shorter distances (<200 kb from the CTCF peaks) but continue for ∼200 kb longer than in wt-CTCF cells (Figure 5F-G; red arrow). Thus, the observed longer stripes in ΔRBR-CTCF mESCs confirm a key prediction of the loop extrusion model and can be explained by loss of the RBR-dependent subset of CTCF boundaries; that is, after an extruding cohesin has reached one CTCF site, it will, on average, need to extrude for longer to reach the next CTCF site, which will result in longer stripes (Figure 5F-G). We also note that CTCF-dependent boundaries are noticeably weaker in ΔRBR-CTCF mESCs (Figure 5F; green arrow), which is consistent with our other results (Figure 4). In summary, our Micro-C analysis reveals that the CTCF RBR domain critically regulates genome organization at the level of TADs, loops, and stripes in mESCs, without affecting A/B compartments.

### Loss of the CTCF RBR reveals distinct sub-classes of TADs and loops

We next asked why some CTCF boundaries depend on the RBR but others do not (Figure 5D). First, we tested whether the RBR is required for CTCF interaction with cohesin using coIPs. Both wt-CTCF and ΔRBR-CTCF immunoprecipitation pulled down cohesin (subunits Rad21 and Smc1a in Figure 6A and Figure S5A). This is especially notable since the protein stability of ΔRBR-CTCF during the IP procedure was significantly reduced (compare CTCF inputs in Figure 6A and S5A). Thus, CTCF interacts with cohesin in an RBR-independent manner, implying that loop loss is not simply due to a failure of ΔRBR-CTCF to interact with cohesin.

Next, we analyzed our CTCF and cohesin (Smc1a) ChIP-Seq data for wt-CTCF and ΔRBR-CTCF mESCs in more detail (Figure 6B; Figure S5B). Consistent with FRAP experiments, which showed no change in residence time at cognate binding sites for ΔRBR-CTCF (Hansen et al., 2018b), ΔRBR-CTCF also still binds the majority of CTCF sites, although the number and occupancy levels were generally reduced (63% of 81,785 wt-CTCF ChIP-Seq peaks maintained in ΔRBR-CTCF mESCs; spike-in normalized ChIP-Seq in Figure S5B). Similarly, about 60% of the cohesin binding sites detected in wt-CTCF mESCs were also occupied in ΔRBR-CTCF mESCs (Figure 6C, Figure S5B). As briefly discussed above, we were surprised to find regions with increased Smc1a binding in ΔRBR-CTCF mESCs compared to wt mESCs (∼40% of all Smc1a peaks in ΔRBR-CTCF mESCs; Figure 6C-D, Figure S5C). The new, “cryptic” cohesin (Smc1a) sites were independent of CTCF and enriched by ∼14-fold at active promoters/chromatin regions compared to other genomic positions (n=2723). Moreover, the cryptic cohesin sites frequently overlapped with binding motifs of master transcription factors in mESCs, such as Oct4/Sox2 and members of the Klf family (Figure 6E). Cryptic cohesin binding was robust enough to stabilize chromatin loops (Figure 6F), and also to increase insulation strength slightly (Figure S5D) (see Methods for details). In light of the loop extrusion model, we speculate that cryptic cohesin binding and loops may arise when mutant ΔRBR-CTCF fails to halt DNA-extruding cohesin, which is then instead halted by bulky transcriptional complexes at nearby transcriptionally active loci. In summary, while we observe ∼17,500 new cohesin (Smc1a) peaks, about 40% of CTCF and cohesin binding sites are lost in ΔRBR-CTCF mESCs, which is in line with our hypothesis that the RBR only regulates a subset of TADs and loops.

To further dissect the site-specific features from the genome-wide average, we divided loops into four quartiles (Figure 6G), such that Q1 contains loops that are largely lost in ΔRBR-CTCF mESCs and Q4 contains loops that are largely unaffected or even strengthened in ΔRBR-CTCF mESCs. We then characterized the CTCF and Smc1a binding profiles at both anchors of loops and only analyzed loops that satisfy three prerequisites: 1) CTCF shows ChIP-Seq signal at both anchors in wt cells; 2) cohesin (Smc1a) shows ChIP-Seq signal at both anchors in wt cells; 3) a pair of convergent CTCF cognate sites are present at both anchors. We then analyzed CTCF and cohesin (Smc1a) ChIP enrichment at the filtered loop anchors for each quartile (Figure 6H). Consistent with a crucial role for CTCF and cohesin, Q1 loops that were disrupted the most in ΔRBR-CTCF mESCs had the lowest CTCF and cohesin occupancy in ΔRBR-CTCF mESCs (see also histograms in Figure S5E), while they were just as strongly, if not more, occupied as Q2-Q4 loops in wt-CTCF mESCs.

In summary, our coIP and ChIP-Seq analyses are consistent with our hypothesis that there are at least two sub-classes of CTCF-sites: an RBR-independent and an RBR-dependent class. Correspondingly, TADs and loops can also be classified as RBR-independent or RBR-dependent. Moreover, our qualitative ChIP-Seq analysis suggests that RBR-dependent loops can be further sub-classified into at least 2 types depending on their CTCF/cohesin dependence (Figure 5D). If this interpretation is correct and robust, we should be able to recover these types naturally after applying an unsupervised clustering algorithm. To test this, we applied *k*-means clustering (using *k*=3) on the most-affected loops (Q1) and recovered 3 loop clusters, similar to Figure 5D (Figure 6I and Figure S5F). Cluster 1 and 2 loops (76%) are lost due to partial and near-complete loss of CTCF/cohesin binding, respectively (type 1 in Figure 5D); cluster 3 loops (24%) are affected loops without strong CTCF/cohesin loss (type 2 in Figure 5D). Thus, this analysis confirms our qualitative assessment in Figure 5D.

Could the CTCF loop type be encoded in the DNA-binding sequence motif? We performed *de novo* motif discovery on the 4 loop quartiles and observed distinct CTCF binding sequence preferences and potential co-regulators (Figure S6A-B). We conclude that loops can be classified into two classes, RBR-dependent and RBR-independent, and that the RBR-dependent class can be further sub-classified into 2 types with distinct CTCF/cohesin binding profiles, and that each class correlates with a distinct CTCF DNA-binding motif preference.

Finally, we asked which other genomic features correlate with RBR-dependent versus RBR-independent loops. We performed an extensive bioinformatics comparison using 70 previously published datasets in mESCs (Figure S6C). Notably, Q4 loops that were not disrupted in ΔRBR-CTCF mESCs correlated with transcriptionally active genomic regions (enhancers, promoters; Figure 6J) and were more frequently found in the A-compartment (Figure S6D), which is generally associated with active genes. In contrast, Q1 loops were relatively larger and more enriched in the B-compartment, which is generally associated with transcriptional repression. These results, albeit inherently correlative, argue against a “*cis*-model” where nascent RNA transcripts stabilize CTCF boundaries in an RBR-dependent manner. Instead, since active sites of transcription are enriched at TAD boundaries (Dixon et al., 2012; Merkenschlager and Nora, 2016), it seems plausible that active transcription may compensate for CTCF boundary weakening in Q4 loops through a CTCF-independent mechanism. In agreement with this hypothesis, despite showing a CTCF ChIP-Seq signal, only a minority of Q4 loop anchors (<40%) contained a canonical CTCF motif, while we could readily detect CTCF binding sites at ∼65% of Q1 loop anchors.

## DISCUSSION

In this report, we have identified unexpected roles for the RNA-binding region (RBR) in CTCF. We confirmed that CTCF self-associates in a largely RNA-mediated manner (Saldaña-Meyer et al., 2014)(Figure 1C) and now demonstrate that the CTCF RBR contributes to CTCF self-association and clustering *in vivo* (Figure 7A). Moreover, we show that the DNA-target search mechanism for CTCF is rendered more efficient by its RBR and describe in detail the RBR-dependent anisotropic diffusion mechanism in a companion paper (Hansen et al., 2018b). Here, we surprisingly find that almost half of all CTCF loops are lost in ΔRBR-CTCF mESCs, suggesting that CTCF-mediated loops can be classified into at least two major classes (Figure 7B): RBR-independent and RBR-dependent CTCF loops. Intriguingly, this may provide a means for differentially engaging or disrupting specific CTCF loops during development and cellular differentiation (Bonev et al., 2017; Pekowska et al., 2018). We discuss some of the implications below.

**Figure 7:**
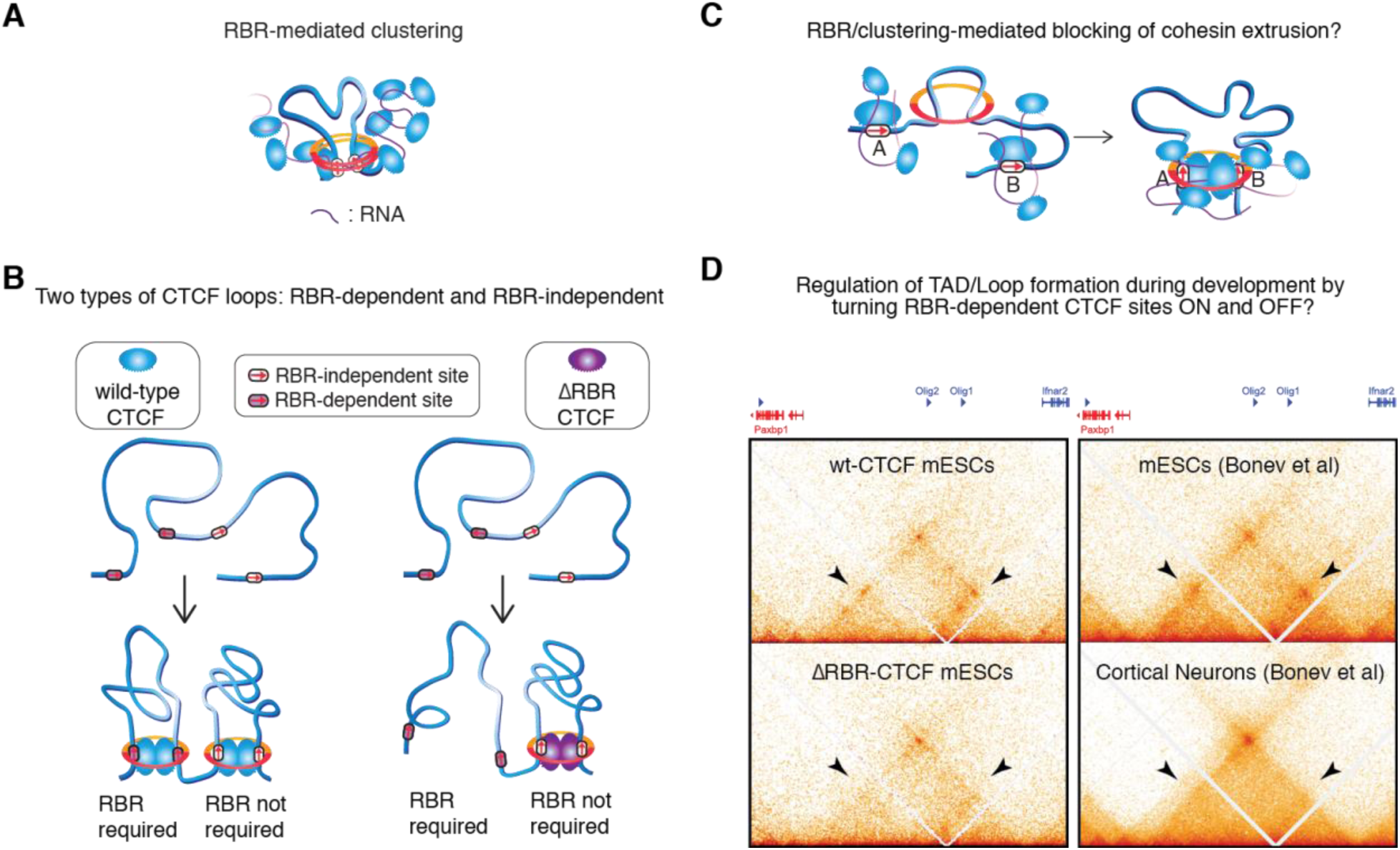
Speculative models for the role of CTCF’s RBR. (**A**) Sketch of a CTCF cluster. Since CTCF clustering is largely RBR-mediated, we speculate that CTCF clusters could be held together by RNA(s), perhaps near loop anchors. (**B**) Two types of CTCF loops. Our analysis of ΔRBR-CTCF mESCs uncovers the existence of at least 2 types of CTCF loops: RBR-dependent and RBR-independent loops. (**C**) Does CTCF clustering help block extruding cohesin? Speculative model that clustering of an otherwise small CTCF protein may contribute to efficiently blocking extruding cohesins. (**D**) Regulation of loops and TADs during differentiation. The ability to turn ON and OFF RBR-dependent CTCF boundaries could potentially provide the means for regulating specific TADs and loops during development by regulating RBR-interaction partners. Here shows a side-by-side comparison of 3D genome reorganization in ΔRBR-CTCF mESCs and differentiated cells at the region around the Olig1 and Olig2 genes (Hi-C data from (Bonev et al., 2017)). Sub-domains and loops (black arrows) are lost in both ΔRBR mESCs and cortical neurons.

### How do CTCF and cohesin interact?

Despite their critical role in 3D genome organization, we know surprisingly little mechanistically about CTCF and cohesin. The function of CTCF’s largely unstructured N- and C-terminal domains remain largely unknown (Martinez and Miranda, 2010; Merkenschlager and Nora, 2016). Similarly, although the related SMC-complex condensin has been observed to extrude loops *in vitro* (Ganji et al., 2018), *in vitro* single-molecule studies of cohesin failed to detect extrusion (Davidson et al., 2016; Kanke et al., 2016; Stigler et al., 2016). Moreover, whether a hypothetical cohesin-based extrusion complex would exist as a single ring or perhaps as a pair of rings remains unclear and a matter of active debate (Cattoglio et al., 2018; Nasmyth, 2011; Sanborn et al., 2015; Skibbens, 2016). Finally, how CTCF and cohesin interact *in vivo* remains to be elucidated. Xiao *et al.* reported that the 575-611 region in human CTCF interacts directly with the SA2-subunit of cohesin and that interaction with the other cohesin subunits is indirect (Xiao et al., 2011). This region largely corresponds to the RBR and is entirely deleted in our ΔRBR-CTCF mESCs. Nevertheless, we observed robust coIP of the cohesin subunits Rad21 and Smc1a with ΔRBR-CTCF (Figure 6A; Figure S5A). Similarly, Saldaña-Meyer *et al.* observed coIP between human ΔRBR-CTCF with the cohesin subunit SA1 (Saldaña-Meyer et al., 2014). Therefore, both our new studies and those of Saldaña-Meyer *et al.* indicate that ΔRBR-CTCF can still interact with cohesin, in apparent contradiction to the work of Xiao *et al.* Can these divergent findings be reconciled? Whereas mammalian cohesin always contains Smc1, Smc3 and Rad21, the SA/STAG/Scc3 subunit can be encoded by either SA1 or SA2. Moreover, mammalian cohesin complexes contain either SA1 or SA2, but not both (Sumara et al., 2000). Curiously, SA2, but not SA1, is among the 12 most frequently mutated proteins in cancer along with well-known onco-proteins such as p53 and Ras (Lawrence et al., 2014).

Furthermore, although cohesin also mediates sister-chromatid cohesion, loss-of-function SA2 mutations in cancer do not cause chromosome segregation defects (Balbás-Martínez et al., 2013). This suggests that it is cohesin’s looping function that is lost in SA2-mutant cancers, and that there are perhaps two functionally distinct cohesin complexes in mammals: SA1-cohesin and SA2-cohesin. Xiao *et al.* observed that CTCF also interacts with SA1-cohesin, though they did not map the relevant region. Whereas SA2 is more abundant than SA1 in somatic cells (Sumara et al., 2000), SA1 and SA2 are expressed roughly equally at the RNA level in our mESCs (Figure S6E). Thus, it is possible that CTCF interacts with SA1-cohesin through another region of the protein while the SA2-cohesin interaction occurs via the RBR. Our coIPs (Figure 6A) cannot rule out that the CTCF-SA2 interaction is disrupted in ΔRBR-CTCF, but that the CTCF-SA1 interaction remains. This could potentially also help explain our observation that CTCF loops fall into RBR-dependent and RBR-independent classes. Thus, it is possible that ΔRBR-CTCF can block SA1-cohesin extrusion, but not SA2-cohesin extrusion, such that SA1-loops are largely unaffected, whereas SA2-loops are lost.

Of course, this raises the question of why some loops would depend on SA1-cohesin and others on SA2-cohesin. CTCF binds DNA through 11 Zinc Fingers (ZFs) and which ZFs contribute to DNA-binding is somewhat idiosyncratic and binding-site dependent (Hashimoto et al., 2017; Nakahashi et al., 2013; Yin et al., 2017). While the core CTCF DNA motif is bound by the central Zinc Fingers (ZFs), only the upstream motif is bound by ZF9-11 (Nakahashi et al., 2013). Since the RBR is just downstream of ZF9-11 (Figure 1A; 2A), it is tempting to speculate that depending on whether or not ZF9-11 are engaged in DNA-binding, there could be allosteric control over which potential RBR interaction partners, RNA(s) or SA2-cohesin, would be engaged. Consistent with this interpretation, we observed clear differences in DNA motifs bound by RBR-dependent and RBR-independent CTCF loops (Figure S6A-B).

We also note that within the context of the loop extrusion model, it is unclear how a ∼3-5 nm sized protein, CTCF, can efficiently block a large and rapidly extruding cohesin complex with a lumen of ∼40-50 nm – and do so in an orientation-specific manner (Guo et al., 2015; Rao et al., 2014; Vietri Rudan et al., 2015; de Wit et al., 2015). We previously showed that CTCF forms clusters in mESCs and U2OS cells (Hansen et al., 2017) and Zirkel *et al.* reported that CTCF forms large foci in senescent cells (Zirkel et al., 2018). Here, we now show that CTCF clustering is partly mediated by the RBR and simultaneously, that the RBR is required for a large subset of loops. It is thus tempting to speculate that cluster and loop formation are related: in particular, RBR-mediated CTCF clustering could make CTCF a more efficient boundary to cohesin-mediated extrusion in at least two ways (Figure 7C): 1) a cluster containing several CTCF proteins, aided by binding to polymers such as RNA, should be much larger and thus more efficient at arresting cohesin than a single chromatin-bound CTCF protein; 2) if CTCF binds cohesin through a specific protein region, having more CTCFs present would increase the probability of a correct encounter between this target interaction surface and cohesin.

While our work here rules out the RBR as the exclusive CTCF region responsible for interacting with cohesin consistent with (Saldaña-Meyer et al., 2014), we suggest that fully elucidating how CTCF and cohesin interact should be an important direction for future research.

### What does the CTCF RBR bind?

We find that CTCF self-association is strongly reduced upon treatment with RNase A *in vitro* (Figure 1C) and that ΔRBR-CTCF shows substantially less clustering in cells (Figure 2C-E). Likewise, others found using fractionation studies that the CTCF RBR is necessary and sufficient for CTCF multimerization *in vitro* (Saldaña-Meyer et al., 2014). Saldaña-Meyer also reported that CTCF directly binds the p53 antisense RNA transcript, Wrap53, and that ZF10-11 contributes to RNA-binding. It is therefore tempting to infer that CTCF clusters are held together by RNA (Figure 7A). However, we emphasize that our results cannot distinguish direct CTCF-RNA binding from a model where the CTCF RBR binds another factor, which then indirectly contributes to CTCF self-association and clustering in an RNase-sensitive manner. Nevertheless, it is worth considering other CTCF-RNA interactions that have been reported beyond Wrap53. CTCF has been reported to directly bind the lincRNA HOTTIP (Wang et al., 2018), the RNA Jpx has been reported to evict CTCF from the X chromosome (Sun et al., 2013), CTCF has been shown to bind RNAs specifically and with high affinity *in vitro* (Kung et al., 2015), and CTCF was also reported to bind the RNA helicase p68/DDX5 together with the noncoding RNA, SRA (Yao et al., 2010). However, there are likely many more CTCF RBR interaction partners and identifying these will be an important but challenging future endeavor.

A recent study identified hundreds of conserved lncRNAs, topological anchor point RNAs (tapRNAs), whose promoters tend to overlap with CTCF loops (Amaral et al., 2018). However, we find that the loops least affected in ΔRBR-CTCF mESCs tend to be associated with active nascent transcription (Figure 6J). While only correlative, we believe this argues against a “*cis* model” where the CTCF RBR predominantly functions by binding nascent RNAs. Does this mean that non-coding RNAs transcribed *in trans* regulate RBR-dependent chromatin loops? This will be an interesting future direction and we note that several non-coding RNAs have been implicated in 3D genome regulation (Melé and Rinn, 2016).

### Regulation of CTCF loops during differentiation and development

An enduring paradox has been the fact that CTCF and cohesin are present in all cell types. Thus, if they were the only factors forming loops and TADs, how can we explain the observation that many TADs and loops change during differentiation (Bonev et al., 2017; Pekowska et al., 2018)? Here we report that CTCF loops can be divided into at least two classes: RBR-dependent and RBR-independent. Moreover, within the RBR-dependent CTCF loop class, we identify at least two types (Figure 5D, 6I and Figure S5F). Having multiple types of CTCF-boundaries provides potential mechanisms through which individual boundaries can be regulated. For example, if CTCF RBR-dependent boundaries function in part by binding other proteins or RNAs, then regulating the abundance or function of these – yet to be identified – factors would provide a potential mechanism for distinct cell types to regulate specific boundaries and CTCF loops during development and differentiation (Figure 7D). Ultimately, this may enable cells to dissolve and form new CTCF-mediated chromatin loops during development and differentiation to regulate enhancer-promoter contacts and establish proper cell-type specific gene expression programs.

### Materials and Methods; Supplementary Figures

Detailed methods and Supplementary Figures and Tables have been uploaded as a separate Supplementary Information file. Key computational tools used include:

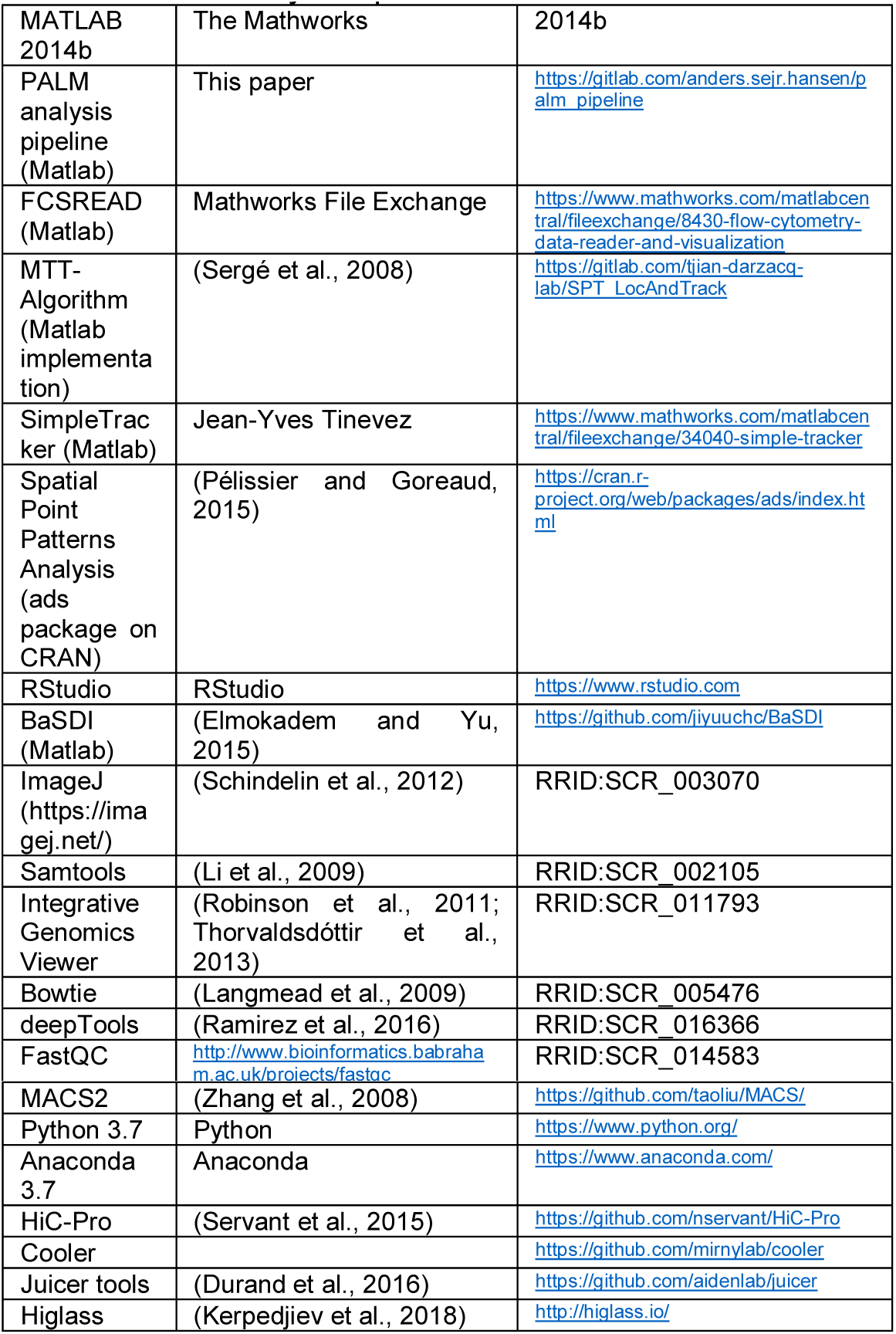

## Supporting information

## Acknowledgements

We thank Luke Lavis for generously providing JF dyes, Hervé Marie-Nelly for help with R-code, Gina M. Dailey for assistance with cloning, Carla J. Inouye for help with the biochemical assays, Assaf Amitai for insightful discussion, Astou Tangara and Ana Robles for microscope assembly and maintenance, Ji Yu and Jean-Yves Tinevez for coding discussions, Daniel J. Lee for help with genotyping, and Dr. Kartoosh Heydari at the Li Ka Shing Facility for flow cytometry assistance. We thank Thomas Graham, and other members of the Tjian and Darzacq labs for comments on the manuscript. This work was performed in part at the CRL Molecular Imaging Center, supported by the Gordon and Betty Moore Foundation. This work used the Vincent J. Coates Genomics Sequencing Laboratory at UC Berkeley, supported by NIH S10 Instrumentation Grants 10RR029668 and S10RR027303. ASH is a postdoctoral fellow of the Siebel Stem Cell Institute. This work was supported by NIH grants UO1-EB021236 and U54-DK107980 (XD), the California Institute of Regenerative Medicine grant LA1-08013 (XD), and by the Howard Hughes Medical Institute (003061, RT). We thank Sheila S Teves for providing the preprint template.

## Notes

#### Summary of Updates

References to computational tools are now included in the main text, instead of just in the Supplementary Methods.

