## Supplementary material for "An RNA-binding region regulates CTCF clustering and chromatin looping"

### SUPPLEMENTARY INFORMATION

#### Cell culture

JM8.N4 mouse embryonic stem cells (Pettitt et al., 2009) (male mESCs; Research Resource Identifier: RRID:CVCL_J962; obtained from the KOMP Repository at UC Davis) were grown and handled as described previously (Hansen et al., 2017). Briefly, mES cells were grown on plates pre-coated with a 0.1% autoclaved gelatin solution (Sigma-Aldrich, St. Louis, MO, G9391) under feeder free conditions in knock-out DMEM with 15% FBS and LIF (full recipe: 500 mL knockout DMEM (ThermoFisher, Waltham, MA, #10829018), 6 mL MEM NEAA (ThermoFisher #11140050), 6 mL GlutaMax (ThermoFisher #35050061), 5 mL Penicillin-streptomycin (ThermoFisher #15140122), 4.6 μL 2-mercapoethanol (Sigma-Aldrich M3148), 90 mL fetal bovine serum (HyClone Logan, UT, FBS SH30910.03 lot #AXJ47554)) and LIF. mES cells were fed by replacing half the medium with fresh medium daily and passaged every two days by trypsinization. Cell lines were pathogen tested (IMPACT II test for mESC C59) as described previously (Hansen et al., 2017). All cell lines will be provided upon request.

#### CRISPR/Cas9-mediated genome editing

Genome-editing was performed as previously described (Hansen et al., 2017). Briefly, we co-transfected cells with a repair plasmid and a plasmid encoding Cas9 and the sgRNA (using 2 μg and 1 μg, respectively, per well in a 6-well plate). The Cas9 plasmid was slightly modified from that distributed from the Zhang lab (Ran et al., 2013): 3xFLAG-SV40NLS-pSpCas9 was expressed from a CBh promoter; the sgRNA was expressed from a U6 promoter; and mVenus was expressed from a PGK promoter. We generally designed 2-4 sgRNAs per knock-in and transfected each (or two of them when necessary) in a separate well. The day of transfection, we pooled all transfected cells and FACS-sorted for transfected cells using the mVenus encoding by the Cas9 plasmid. For edits where there was no tag added (e.g. to replace the RBR with 3xHA), we immediately plated single clones after the FACS. But for knock-ins with tags, e.g. 3xFLAG-Halo-CTCF or V5-$SNAP_f$-CTCF, we first grew up cells and then labeled cells with dye (Halo-TMR for 3xFLAG-Halo-CTCF; SNAP-JF646 for V5-$SNAP_f$-CTCF) and then did a second round of FACS-sorting to increase the efficiency. Selected cells were plated at very low density (~0.1 cells per $mm^2$), and single colonies were then picked, expanded and genotyped by PCR. Successfully edited clones were further verified by PCR followed by Sanger sequencing and Western blotting.

For knock-ins with 2 different tags, we generated them using the above protocol in 2 steps. We first isolated a heterozygous knock-in clone for one tag and we then re-edited that clone to introduce the second tag. This was the case for C62, where one of the diploid CTCF alleles is V5-$SNAP_f$-tagged and the other 3xFLAG-Halo-tagged. We first isolated a clone with a correct V5-$SNAP_f$-tagged CTCF allele and a null CTCF allele, where non-homologous end joining event following Cas9 cleavage introduced a 4-nucleotide deletion (81_84delACGC), leading to a premature stop codon. We then designed sgRNAs specific for the null CTCF allele and re-targeted this clone with a 3xFLAG-Halo-CTCF repair vector. To build the repair vectors, we modified a pUC57 plasmid to contain the tag of interest flanked by ~500 bp of genomic homology sequence on either side (IDT gBLocks). To prevent the Cas9-sgRNA complex from cutting the repair vector, we introduced synonymous mutations in the first nine codons after the ATG. To link the SNAP and Halo proteins to CTCF, we used the Sheff and Thorn linker (GDGAGLIN) (Sheff and Thorn, 2004) and a TEV linker sequence (EDLYFQS), respectively. mESC clones were screened using a three-primer PCR (two genomic primers external to the left and right homology sequences, and one internal to the tag).

To endogenously and homozygously delete the RBR region in the previously published C59 mESC line (Hansen et al., 2017), we generated by Gibson Assembly a repair vector modifying a pBlueScript II SK (+) plasmid to contain the Sheff and Thorn linker followed by a 3xHA tag (Figure S1F), and flanked by ~500 bp of genomic homology sequence on either side. mESC clones were screened using a three-primer PCR (one genomic primer external to the left homology sequence, one internal to the right homology region, and an internal HA primer). Notably, we failed to generate clones with a simple deletion of the RBR, possibly because shortening of the already small exon 10 (only 135 bp-long, 27 bp upon RBR deletion) causes exon skipping and aberrant splicing.

All plasmids used in the editing are available upon request as are any of the cell lines. See Table S1 for sgRNA and primer sequences.

#### Cell Cycle phase analysis

Cell cycle phase analysis was performed using the Click-iT EdU Alexa Fluor 488 Flow Cytometry Assay Kit (ThermoFisher Scientific Cat. # C10425) according to manufacturer's instructions, but with minor modifications. C59 mESCs (Halo-CTCF; Rad21-SNAP$_f$) and C59D2 mESCs (ΔRBR-Halo-CTCF; Rad21-SNAP$_f$) were grown overnight in a 6-well plate and labelled with 10 μM EdU for 30 min at 37°C/5.5% $CO_2$ in a TC incubator (one well was unlabeled, as a negative control). Cell were harvested, washed with 1% BSA in PBS, permeabilized (using 100 μL 1x Click-iT saponin-based permeabilization and wash reagent (Component D; see kit manual), mixed well and then incubated for 15 min. 0.5 mL Click-iT reaction was added to each tube and incubated for 30 min in the dark. Cells were washed with 1x Click-iT saponin-based permeabilization and wash reagent and resuspended in 1x Click-iT saponin-based permeabilization and wash reagent with DAPI (5 ng/mL) and incubated for 10 min. Cells were then spun down and re-suspended in 1% BSA in PBS and FACS performed on a LSR Fortessa Cytometer. DAPI fluorescence was excited using a 405 nm laser and collected using a 450/50 bandpass emission filter. Alexa Flour 488 fluorescence was excited using a 488 nm laser and collected using a 525/50 bandpass emission filter. Cells were sorted based on forward and side scattering using identical settings for C59 and C59D2 mESCs. Cell cycle analysis was performed using custom-written MATLAB code using identical settings for C59 and C59D2 mESCs as illustrated in Figure S3A. Three independent biological replicates were performed.

#### CTCF FACS abundance quantification

FACS was performed as previously described (Hansen et al., 2017). We grew C59 mESCs (Halo-CTCF; Rad21-SNAP$_f$) and C59D2 mESCs (ΔRBR-Halo-CTCF; Rad21-SNAP$_f$) overnight in a 6-well plate and labeled 1 well with 500 nM Halo-TMR (Promega Cat. # G8521) and left 1 well unlabeled (negative control for baseline fluorescence). Cells were labeled for 30 min at 37°C/5.5% $CO_2$ in a TC incubator, washed with PBS and incubated with medium for 5 min in a TC incubator. Cells were then washed again with PBS, harvested, filtered and fluorescence quantified in live cells on a LSR Fortessa Cytometer, exciting fluorescence with a 561 nm laser and collecting fluorescence through a 610/20 bandpass emission filter. Live cells were gated based on forward and side scattering (using identical settings for C59 and C59D2 mESCs) using custom-written MATLAB code and the relative abundance quantified as the relative background-subtracted mean fluorescence as illustrated in Figure 2C and S1G.

#### Growth Assay

When passaging cells, two processes contribute to the apparent growth rate: 1) the fraction of cells that survive passaging and 2) the growth rate. To compare exclusively the growth rate of mESC C59 Halo-CTCF and mESC C59D2 ΔRBR-Halo-CTCF, we therefore took the following approach. On day 0, we plated 250,000 cells in 2 wells in a 6-well plate. On day 1, we collected and counted the number of cells from 1 well. This gave us the number of cells that survived plating. Let this number be $N_1$. On day 2, we then collected and counted the number of cells from the second well. Let this number be $N_2$ and the time between the measurements be $\Delta\tau$. The doubling time is then given by:

$$\tau_{\text{DOUBLING}} = \frac{\Delta\tau \ln(2)}{\ln\left(\frac{N_2}{N_1}\right)}$$

We performed 4 biological replicates and grew C59 and C59D2 side-by-side at the same time and handled them identically. The bar-graph in Figure 2D shows the mean and standard error of the mean from the 4 replicates.

#### PALM

PALM was performed as previously described (Hansen et al., 2017) but with minor modifications. C59 mESCs (Halo-CTCF; Rad21-SNAP$_f$) and C59D2 mESCs (ΔRBR-Halo-CTCF; Rad21-SNAP$_f$) were grown overnight on MatriGel coated plasma-cleaned 25 mm circular no 1.5H cover glasses (Marienfeld, Germany, High-Precision 0117650), labelled with 500 nM PA-JF549 (Grimm et al., 2016) for 30 min at 37°C/5.5% $CO_2$ in a TC incubator, washed twice (medium removed; PBS wash; fresh medium for 5 min), and then fixed in 4% Formaldehyde / 0.2% Glutaraldehyde in PBS for 20 min at 37°C, washed with PBS and then imaged in PBS with 0.01% (w/v) $NaN_3$ on the same day. All PALM movies were acquired at room temperature using continuous HiLo illumination on the same microscope as previously described (Hansen et al., 2017). We used the following laser lines: main excitation laser (561 nm for PA-JF549) and photo-activation laser (405 nm). However, the intensity of the 405 nm laser was gradually increased over the course of the illumination sequence to image all molecules and at the same time avoid too many molecules being activated at any given frame. The following camera settings were used: 25 ms exposure time; frame transfer mode; vertical shift speed: 0.9 µs; ROI: variable. In total, 40,000 frames were recorded for each cell (~20 min), which was sufficient to image and bleach all labeled molecules at an effective pixel size of 106.67 nm, which resulted in a mean localization error (defined as the standard deviation) of ~13-14 nm (Figure S1H). We recorded 6-10 movies per cell line per day (and always imaged both C59 and C59D2 on the same day) and performed 3 biological replicates. Each movie contained several nuclei (generally 3-6), which improved the robustness of the algorithmic drift-correction (Elmokadem and Yu, 2015). We obtained and analyzed a total of 52 cells for C59 and 46 cells for C59D2.

Molecules in PALM data were localized using a custom-written Matlab implementation of the MTT-algorithm ((Sergé et al., 2008); code is available on GitLab: https://gitlab.com/tjian-darzacq-lab/SPT_LocAndTrack) and the following settings: Localization error: $10^{-6}$; deflation loops: 0. After localization, the data was analyzed as described below using code available on GitLab: https://gitlab.com/anders.sejr.hansen/palm_pipeline

#### PALM analysis

Full details on PALM analysis as well as code to reproduce our results are available on GitLab: https://gitlab.com/anders.sejr.hansen/palm_pipeline. Here we summarize the major steps. First, drift-correction and merging of blinks is achieved through the main script "DriftCorrectMergeBlinks.m", which calls a number of functions and runs in parallel as default, so the parallel processing toolbox i0n Matlab is neccesary. Drift-correction is first performed using a custom-modified implementation of BaSDI (Elmokadem and Yu, 2015) ("BaSDI_ASH"). This is achieved through the function "IterativeBaSDI_DriftCorrect.m" using FramesBin=2000; PixelBin=10; Iterations=5. Compared with BaSDI, the main difference is that we found multiple iterations to be necessary to reach convergence and we have therefore custom-written the wrapper "IterativeBaSDI_DriftCorrect.m" to achieve this. Since the inferred drift is binned according to "FramesBin", we use linear interpolation to drift-correct each frame. Once drift-correction has been achieved, we merge photo-blinking using a custom implementation of SimpleTracker (https://www.mathworks.com/matlabcentral/fileexchange/34040-simple-tracker), which was modified to be substantially more memory-efficient for large PALM movies (SimpleTracker_ASH). An important aspect of PALM, especially with very photo-stable dyes such as PA-JF549 (Grimm et al., 2016), is that the same molecule can appear in multiple adjacent frames and also blink such that there are gaps. It is therefore essential to link these appearances, which we accomplish using SimpleTracker's implementation of nearest neighbor tracking and we allow a maximal linking distance of 75 nm and maximally 2 gaps. We note that 75 nm is quite lenient since the localization error is less than 15 nm, but we chose it so to ensure we fully correct for multiple appearances. For each molecule with multiple appearances, we collapse all the localizations to a single localization and take the x,y coordinates to be the means.

After drift-correcting and merging, individual nuclei are segmented after Gaussian smoothing of reconstructed images using a 60 nm pixel size. Since the movies contain several nuclei, each nucleus is manually segmented using polygon-segmentation. For each nucleus, a series of summary statistics are then displayed and saved (e.g. localization error, number of localization per frame, nuclear reconstructions) and each nucleus is saved to a separate directory together with code for running K-Ripley analysis (Besag, 1977; Boehning et al., 2018; Ripley, 1976) using the ads package in R (Pélissier and Goreaud, 2015) as well as code for running a Bayesian cluster identification algorithm (Rubin-Delanchy et al., 2015).

The R-code for running K-Ripley analysis was written by Herve Marie-Nelly and is described elsewhere (Boehning et al., 2018). The version included here is a slightly modified version and we refer the reader to the tutorial on GitLab for how to run it (requires both Python and R).

Finally, the results of the K-Ripley analysis were plotted with "PLOT_K_L_g_Ripley.m" and Figure 2G show the mean and standard error of the mean across the population.

The example reconstructions of CTCF nuclear localization in Figure 2E-F were plotted using ViSP (Beheiry and Dahan, 2013). Each molecule was plotted using 25 nm (FWHM) and colored according to the neighbor density (0-200 Neighbors (min/max); Neighborhood Radius: 100 nm; Jet colormap (cMin-cMax: 0-0.35) with identical settings for C59 Halo-CTCF and C59D2 ΔRBR-Halo-CTCF.

#### Western Blotting

Cells were grown in 6-well plates to confluency, washed twice with ice-cold PBS with protease inhibitors and scraped in 300 μl of high salt lysis buffer (0.5 M NaCl, 25 mM HEPES, 1 mM $MgCl_2$, 0.2 mM EDTA, 0.5% NP-40 and protease inhibitors). Lysates were immediately transferred to 1.5 ml tubes containing 100 μl of 4X protein loading buffer (16% 2-Mercaptoethanol, 200 mM Tris-HCl pH 6.8, 8% SDS, 40% glycerol, 400 mM DTT, 0.4% bromophenol blue), boiled for 20' and loaded to 8% Bis-Tris protein gels (10 μl per lane). Proteins were transferred onto nitrocellulose membranes (Amershan Protran 0.45 um NC, GE Healthcare) for 2 hrs at 100V. Membranes were blocked in TBS-Tween with 10% milk for at least 1 hr at room temperature and blotted with the specified antibodies in TBS-T with 5% milk at 4 ˚C overnight. HRP-conjugated secondary antibodies were diluted 1:5000 in TBS-T with 5% milk and incubated at room temperature for an hour prior to the chemiluminescence reaction. Band intensities were measured with the ImageJ "Analyze Gels" function (Schindelin et al., 2012) and used to calculate IP and CoIP efficiencies.

#### Co-immunoprecipitation (CoIP) assays

For CoIP experiments, cells were scraped from plates in ice-cold phosphate-buffered saline (PBS) with PMSF and aprotinin, pelleted, and flash-frozen in liquid nitrogen. Cell pellets where thawed on ice, resuspended to 1 ml/10 cm plate of cell lysis buffer (5 mM PIPES pH 8.0, 85 mM KCl, 0.5% NP-40 and protease inhibitors), and incubated on ice for 10'. Nuclei were pelleted in a tabletop centrifuge at 4 °C, at 4000 rpm for 10', and resuspended to 0.5 ml/10 cm plate of low salt lysis buffer either with or without benzonase (600U/ml) and rocked 4 hrs at 4 °C. After the incubation the salt concentration was adjusted to 0.2 M NaCl final and the lysates were incubated for another 30' at 4 °C. 50 μl of each lysate were used for DNA and RNA extraction (see below), while the rest was cleared by centrifugation at maximum speed at 4 °C and the supernatants quantified by Bradford. In a typical CoIP experiment, 1 mg of proteins was diluted in 1 ml CoIP buffer (0.2 M NaCl, 25 mM Hepes, 1 mM MgCl2, 0.2 mM EDTA, 0.5% NP-40 and protease inhibitors) and pre-cleared for 2 hrs at 4 °C with protein-A/G sepharose beads (GE Healthcare Life Sciences) before overnight immunoprecipitation with 4 mg of either normal serum IgGs or specific antibodies. Some pre-cleared lysate was kept at 4 °C overnight as input. Protein-A/G-sepharose beads precleared overnight in CoIP buffer with 0.5% BSA were then added to the samples and incubated at 4 °C for 2 hrs. Beads were then washed extensively with CoIP buffer, and proteins were eluted by boiling the beads for 5' in 2X SDS-loading buffer, followed by SDS-PAGE and Western Blotting.

#### CoIP DNA and RNA extraction and quantification

For DNA extraction, 50 μl of lysates were added to 150 μl of CoIP buffer and extracted twice with 200 μl of phenol-chloroform (UltraPure™ Phenol:Chloroform:Isoamyl Alcohol (25:24:1, v/v)). After centrifugation at room temperature and maximum speed for 5', the aqueous phase containing DNA was added of 2 volumes of 100% ethanol and precipitated 30' at -80 °C. After centrifugation at 4 °C for 20' at maximum speed, DNA was re-dissolved in 25 μl water and quantified by nanodrop. About 100 ng of the untreated sample DNA, or an equal volume from the nuclease treated samples, were used for relative quantification by quantitative PCR (qPCR) with SYBR Select Master Mix for CFX (Applied Biosystems, ThermoFisher) on a BIO-RAD CFX Real-time PCR system (primer sequences in Table S1).

RNA was extracted from 50 μl of lysates with 500 μl of TRIzol™ reagent, following manufacturer's instructions. The RNA pellet was re-dissolved in 25 μl of water and quantified by nanodrop. About 1 μg of the untreated sample RNA, or an equal volume from the nuclease treated samples, was retrotranscribed with SuperScript™ III Reverse Transcriptase and random examers. cDNA was diluted 1:20 and 2 μl quantified by qPCR as above.

#### Chromatin immunoprecipitation (ChIP)

Smc1a, CTCF and control IgG ChIP assays were performed in the parental C59 ES cell line (wt-CTCF) and in its derivative clone C59D2 (ΔRBR-CTCF). Cells were cross-linked for 5' at room temperature with 1% formaldehyde-containing Knockout D-MEM; cross-linking was stopped by PBS-glycine (0.125 M final). Cells were washed twice with ice-cold PBS, scraped, centrifuged for 10' at 4000 rpm and flash-frozen in liquid nitrogen. Cell pellets were thawed in ice, resuspended in cell lysis buffer (5 mM PIPES, pH 8.0, 85 mM KCl, and 0.5% NP-40, 1 ml/15 cm plate) and incubated for 10' on ice. During the incubation, the lysates were repeatedly pipetted up and down every 5 minutes. Lysates were then centrifuged for 10' at 4000 rpm. Nuclear pellets were measured and resuspended in 6 volumes of sonication buffer (50 mM Tris-HCl, pH 8.1, 10 mM EDTA pH 8.0, 0.1% SDS), incubated on ice for 10', and sonicated to obtain DNA fragments below 2000 bp in length (Covaris S220 sonicator, 20% Duty factor, 200 cycles/burst, 150 peak incident power, 30-40 cycles of 20" on and 40" off). Sonicated lysates were cleared by centrifugation (20' at 13200 rpm) and 625-800 μg of chromatin were diluted in RIPA buffer (10 mM Tris-HCl, pH 8.0, 1 mM EDTA pH 8.0, 0.5 mM EGTA, 1% Triton X-100, 0.1% SDS, 0.1% Na-deoxycholate, 140 mM NaCl) to a final concentration of 0.8 μg/μL, precleared with Protein A sepharose (GE Healthcare) for 2 hrs at 4 °C and immunoprecipitated overnight with 6.25-8 μg of normal mouse IgGs (ChromPure rabbit normal IgG; Jackson ImmunoResearch), anti-Smc1a (Abcam ab154769) or anti-CTCF antibodies (Abcam ab128873). 4% of the precleared chromatin was saved as input. After the overnight incubation, samples were added of 20 μl of Protein A sepharose beads precleared overnight in RIPA buffer with 0.5% (w/v) BSA and incubated for 2 hrs at 4 °C. Immunoprecipitated samples were washed 5 times with RIPA buffer, once with LiCl buffer (0.5% NP-40, 0.5% Na-deoxicholate, 250 mM LiCl, 1 mM EDTA pH 8.0), and once with TE. After the last wash, immunoprecipitated complexes were eluted from the beads twice with 150 μl of TE with 1% SDS, each time incubating 30' in a thermomixer set at 37 °C and 900 rpm. The 300 μl eluted material was added of 1 μl of RNaseA (10 mg/ml) and 18 μl 5M NaCl, and incubated at 67 °C for 4-5 hrs to reverse formaldehyde cross-linking. To inputs were added elution buffer to 300 μl total volume, and subject to the same treatment. Reverse cross-linked samples were added of 2.5 volumes of ice-cold ethanol and precipitated overnight at -20 °C. DNA was pelleted by centrifugation (20' at 13,200 rpm and 4 °C), and pellets resuspended in 100 μl TE, 25 μl 5X PK buffer (50 mM Tris-HCl, pH 7.5, 25 mM EDTA pH 8.0, 1.25% SDS), and 1.5 μl of proteinase K (20 mg/ml), and incubated 2 hrs at 45 °C. After proteinase K digestion, DNA was purified with the Qiagen QIAquick PCR Purification Kit, eluted in 60 μl of water and used for ChIP-Seq library preparation as described below.

#### ChIP-Seq library preparation

ChIP-Seq libraries were prepared independently from two ChIP biological replicates using the Solexa rapid library protocol. Briefly, immunoprecipitated DNA or 50 ng of input DNA was end-repaired, phosphorylated and adenylated in a single 50 μl reaction containing 31.5 μl of DNA, 5 μl of spike-in yeast DNA from MNase treated nucleosomes (10 ng/ml) (Skene and Henikoff, 2017) and 13.5 μl of end-repair/3' A mix. Reactions were incubated in a thermal cycler for 15' at 12 °C, 15' at 37 °C, 20' at 72 °C, and held at 4 °C.

| End-repair/3' A mix component | Final concentration | Cat # |
|---|---|---|
| 10X T4 DNA ligase buffer | 1X | NEB #B0202S |
| 10 mM dNTPs | 0.5 mM each | KAPA #KK1017 |
| 10 mM ATP | 0.25 mM | NEB #P0756S |
| 40% PEG 4000 | 2.5% | |
| 10 U/μl T4 PNK | 0.0025 U/μL | NEB #M0201S |
| 5U/μl T4 DNA polymerase* | 0.0025 U/μL | Invitrogen #18005025 |
| 5U/μl Taq DNA polymerase** | 0.0025 U/μL | Thermo #EP0401 |

* diluted 1:20 in 1x T4 DNA

ligase buffer
** diluted 1:20 in 1X standard Taq buffer (NEB #B9014S)

To reactions were added 4 μl of water, 1 μl of Illumina TruSeq adapters, 55 μl of 2x Rapid DNA ligase buffer (Enzymatics #B101L) and 5 μl of DNA ligase (Enzymatics #L6030-HC-L), and incubated for 15' at 20 °C. Ligations were cleaned up twice with AMPure XP beads (Agencourt #A63880) diluted 1:2 with 20% PEG, 1.25M NaCl (first cleanup: 38 μl; beads eluted with 53 μl of 10 mM Tris-HCl pH 8.0, 50 μl transferred to a new tube and added of 55 μl of beads:PEG solution). Final elution volume was in 22 μl of 10 mM Tris-HCl pH 8.0, 20 μl of which were transferred to a new tube and amplified by PCR (45" at 98 °C; 14 cycles of 15" at 98 °C and 10" at 60 °C; 1' at 72 °C; hold at 4 °C).

| PCR mix component | Final concentration | Cat # |
|---|---|---|
| 5X KAPA buffer | 1X | KAPA #KK2502 |
| 10 mM dNTPs | 0.3 mM each | KAPA #KK1017 |
| 5μM TruSeq PCR primers | 0.5 μM | Primer sequence in Table S1X |
| KAPA HS HIFI polymerase | 1 U | KAPA #KK2502 |
| Nuclease-free water to 30 μL | | |

PCR reactions were cleaned up once with 38 μl of AMPure XP beads diluted 1:2 with 20% PEG, 1.25M NaCl and eluted with 33 μl of 10 mM Tris-HCl pH 8.0, 30 μl of which were transferred to a new tube. We assessed library quality and fragment size by qPCR and Fragment analyzer™, and sequenced 8-12 multiplexed libraries per lane on the Illumina HiSeq4000 sequencing platform (single end-reads, 50 bp long) at the Vincent J. Coates Genomics Sequencing Laboratory at UC Berkeley (supported by NIH S10 OD018174 Instrumentation Grant).

#### ChIP-Seq analysis

Input, IgG, Smc1a and CTCF ChIP-Seq raw reads from wt-CTCF (C59) and ΔRBR-CTCF (C59D2) ESCs (16 libraries total) were quality-checked with FastQC (http://www.bioinformatics.babraham.ac.uk/projects/fastqc) and aligned onto the mouse and the yeast genome (mm10 and sacCer3 assembly, respectively) using Bowtie (Langmead et al., 2009), allowing for two mismatches (-n 2) and no multiple alignments (-m 1). We used Samtools ((Li et al., 2009) version 1.9) to sort and index bowtie output .bam files, remove duplicates from mapped reads (rmdup -s) and merge ChIP-Seq replicates. Peaks were called with MACS2 (--nomodel --extsize 300) (Zhang et al., 2008) using input DNA as a control. Overlap between ChIP-Seq peaks across samples were computed through Galaxy (Blankenberg et al., 2010; Giardine et al., 2005; Goecks et al., 2010), requiring a minimum 1-bp overlap between peak intervals.

For spike-in control normalization, we performed pairwise comparisons (*e.g.*, C59 input *vs.* C59D2 input) and selected the sample with the lowest number of unique yeast alignments (C59D2 input: 94,642 reads *vs.* C59 input: 119,846 reads). We then used this value to compute a scale factor (*sf*) for the other sample (C59 *sf*: 94,642 / 119,846 = 0.79), to be used in the downstream analyses (see below).

To create heatmaps we used deepTools (version 2.4.1) (Ramirez et al., 2016). We first ran bamCoverage (--binSize 50 --extendReads 300 -of bigwig) and normalized read numbers to either 1x sequencing depth (--normalizeTo1x 2150570000) or to the spike-in yeast DNA (--scaleFactor *sf*), obtaining read coverage per 50-bp bins across the whole genome (bigWig files). We then used the bigWig files to compute read numbers across 6 kb centered on C59 CTCF or Smc1a peak summits as called by MACS2 (computeMatrix reference-point --referencePoint=TSS --upstream 3000 --downstream 3000 --missingDataAsZero --sortRegions=no). We sorted the output matrices by decreasing C59D2 enrichment, calculated as the total number of reads within a MACS2 called ChIP-Seq peak. Finally, heatmaps were created with the plotHeatmap tool (--averageTypeSummaryPlot=mean --colorMap='Blues' --sortRegions=no).

Enriched regions were visualized on the mm10 genome with the Integrative Genomics Viewer (IGV) (Robinson et al., 2011; Thorvaldsdóttir et al., 2013) using the bigWig output files from deepTools bamCoverage.

#### Micro-C

Mammalian Micro-C protocol and analysis were modified from (Hsieh et al., 2016). Here, we briefly summarize the key concepts of Micro-C experiment and data analysis. The detailed step-by-step protocol can be found in

Supplemental Protocol.

**I. Prepare crosslinked chromatin from cell culture**

One to five million of trypsinized mouse embryonic stem cells were directly resuspended and crosslinked with freshly made 1% formaldehyde at room temperature for 10 minutes. Crosslinking reaction was quenched by adding Tris buffer (pH=7.5) to final 0.75M at room temperature. Crosslinked cells were washed twice by 1x PBS and subjected to the second crosslinking with 3mM DSG crosslinking solution for 45 minutes at room temperature. Cells were snap-frozen and can be stored at -80°C up to a year.

**II. Digest crosslinked chromatin by micrococcal nuclease**

Crosslinked cells were permeabilized in ice-cold Micro-C Buffer #1 (50mM NaCl, 10mM Tris-HCl pH=7.5, 5mM MgCl2, 1M CaCl2, 0.2% NP-40, 1x Protease Inhibitor Cocktail) for 20 minutes. Chromatin from permeabilized cells was digested by pre-titrated concentration of micrococcal nuclease to about 90% of mononucleosomes and 10% dinucleosome at 37°C for 10 minutes. Digestion reaction was stopped by adding EGTA to a final concentration at 4mM and incubated at 65°C for 10 minutes to completely deactivate enzyme activity. MNase-digested chromatin was washed twice with ice-cold Micro-C Buffer #2 (50mM NaCl, 10mM Tris-HCl pH=7.5, 10mM MgCl2).

**III. Repair fragment ends**

Digested chromatin fragments were then subjected to dephosphorylation, phosphorylation, end-chewing reactions by T4 Polynucleotide Kinase and DNA Polymerase I Klenow Fragment in Micro-C end-repair buffers (50mM NaCl, 10mM Tris-HCl pH=7.5, 10mM MgCl2, 100ug/mL BSA, 2mM ATP, 5mM DTT, no dNTPs) at 37°C for 30 minutes. Blunt-end reaction was triggered by adding biotin-dATP, biotin-dCTP, dGTP, and dTTP to a final concentration at 66mM and incubated at 25°C for 45 minutes. Fill-in reaction was stopped by 30mM EDTA in 65°C for 20 minutes. Chromatin was washed once with ice-cold Micro-C Buffer #3 (50mM Tris-HCl pH=7.5, 10mM MgCl2).

**IV. Proximity ligation and purge biotin-dNTP from unligated ends**

Chromatin fragments with biotin-dNTPs were then ligated by T4 DNA ligase at room temperature for at least 2 hours. Unligated ends containing biotin-dNTPs were then removed by 5' to 3' exonuclease III in 37°C for at least 15 minutes. Chromatin was subjected to reverse crosslinking and protein digestion in proteinase K buffer (2mg/mL Proteinase K, 1% SDS, 1x TE buffer) in 65°C overnight.

**V. Purify dinucleosomal DNA**

DNA from Micro-C sample was extracted by Phenol:Chloroform:Isoamyl Alcohol (25:24:1) solution and ethanol precipitation method. DNA was cleaned-up by Zymo DNA clean and concentration column prior to gel size-selection for dinucleosomal DNA at 200 to 400 bp. DNA was then extracted from agarose gel by Zymo Gel purification column.

**VI. Micro-C library preparation for deep sequencing**

Purified DNA with biotin-dNTPs was captured by Dynabeads® MyOne™ Streptavidin C1. Standard library preparation protocol including end-repair, A-tailing, and adapter ligation was performed on beads with individual enzymes purchased from Lucigen or NEBnext UltraII kit. Sequencing library was amplified by Kapa HiFi PCR enzyme with lowest possible cycles to reduce PCR duplicates. Sequencing library was sequenced paired-end 50x50 or 100x100 in Illumina HiSeq 4000 sequencer.

**VII. Micro-C data analysis**

**a. Mapping and pairing Micro-C contacts.**

Micro-C data were preprocessed by a streamlined pipeline HiC-pro (Servant et al., 2015). Briefly, sequencing reads were mapped to mouse mm10 genome by Bowtie2 in –very-sensitive-local mode. Mapped reads were paired and pairs with multiple hits, low MAPQ, self-circle, and PCR duplicates were removed. Output file containing all valid pairs were used for following analysis.

**b. Visualize Micro-C data.**

Micro-C data was converted to standard 4DN formats (e.g. .cool or .hic file) with multiple resolutions, typically ranging from 500bp to 10Mb. Cool file can be visualized on Higlass browser (http://higlass.io/) and hic file can be visualized on Juicebox (https://github.com/aidenlab/Juicebox). All files can be found at https://www.ncbi.nlm.nih.gov/geo/query/acc.cgi?acc=GSE123636. In this study, all snapshots of contact matrices were generated by Higlass browser.

c. **Binning and balancing of Micro-C data.**
Valid pairs were binned to 500bp resolution by using cooler packages (https://github.com/mirnylab/cooler) for cool file or juicer packages (https://github.com/aidenlab/juicer) for hic files. Low mappability or noisy regions were precluded prior to matrix balancing. Matrix was balanced by using iterative correction (IC) for cool file or Knight-Ruiz (KR) for hic file. Visually, both normalization methods generate equal quality of contact maps. Multiple resolutions of contact maps can be generated by using matrix coarsen or zoomify functions in cooler package.

d. **Contact probability analysis**
Only intra-chromosomal contacts were used to calculated contact density in bins with exponentially increasing widths from 100bp to 10Mb. Contacts shorter than 100bp were removed to minimize noises introduced by self-ligation or unligated products. The numbers of intra-chromosomal contacts within the range of distance were calculated and normalized by the all contacts within this range. Plot shown in Figure 3D only included pairs in "UNI" direction to minimize noise from unligated products.

e. **Compartment analysis**
Chromosomal compartments were identified using principal component analysis (PCA) on contact maps in 100kb resolution. The first component typically represents the compartment profile in mammalian genome – positive eigenvector value enriches with A compartment (gene-rich regions) and negative eigenvalue enriches with B compartment (gene-poor regions). The rank of compartment strength shown in saddle plot was analyzed by rearrangement and aggregation of genome-wide distance-normalized contact matrix in the order of increasing eigenvector values.

f. **Domain/boundary analysis**
We used two approaches to identify TADs and TAD boundaries/insulators. The detailed methods were described in (Crane et al., 2015) for the insulation score analysis or (Rao et al., 2014) for the arrowhead transformation analysis. Briefly, the optimal condition for calling insulation score was determined by testing multiple sizes of sliding window on 10kb - 100kb resolutions of contact maps. A sliding window at 200kb on 20kb-binned contact maps was used to analyze insulation score in this study. The signal within the sliding window was assigned to the corresponding bin across the entire genome. The insulation score was normalized by calculating the log2 ratio of individual score and the mean of genome-wide averaged insulation score. TAD boundaries/insulators can be identified by calling the local minima along the normalized insulation score. The arrowhead analysis defined as $A_{i,i+d} = (M^*_{i,i-d} - M^*_{i,i+d})/(M^*_{i,i-d} + M^*_{i,i+d})$. $A_{i,i+d}$ can be thought of as the measurement of the directionality preference of locus $i$, restricted to contacts at a linear distance of $d$. $A_{i,i+d}$ will be strong positive / negative if either one of $i,i-d$ or $i,i+d$ is inside the domain and one another is not, but $A_{i,i+d}$ will be close to zero if both loci are inside or outside the domain. Assigning this query across the genome, the edges of domain will be sharpened and TADs can be detected. For aggregate domain analysis, the individual TADs were rescaled to the same size as $TAD_{i,j}=((C_i\text{-}TAD_{start})/(TAD_{end}\text{-}TAD_{start}), (C_j\text{-}TAD_{end})/TAD_{end}\text{-}TAD_{start}))$, where $C_{i,j}$ is a pair of contact loci within a TAD. The rescale matrices were then aggregated at the center of plot with either ICE or distance normalization.

g. **Loop analysis**
Loops in mouse ES cells were discovered by HiCCUPS as described in (Rao et al., 2014) HICCUPS uses a modified Benjamini-Hochberg FDR control procedure to reduce the rate of false positive and identify highly reliable loop annotations on contact map. Loops were called on multiple resolutions (1, 5, and 10kb) of KR-normalized Micro-C contact matrices with a false discovery rate smaller than 0.1. Peak widths and windows of peak-to-merging were set as 5kb for 1kb contact maps and 20kb for 5kb and 10kb contact maps. Genome-wide loop comparison/quantification was assessed by using aggregate peak analysis. All called loops were compiled on a center of 25kb x 25kb matrix with 1kb resolution of KR-normalized data. Loops within 55kb of diagonal were excluded to avoid distance decay effects. The ratio of loop enrichment was calculated by dividing observed contact in a searching window by the expected bottom-left submatrix.

h. **Pile-up analysis**

The concept of pile-up analysis is similar to aggregate domain analysis described above. Briefly, we used a set of ChIP-Seq peaks of interest (e.g. CTCF ChIP-Seq in this study) as baits to extract 600kb x 600kb snippets of contact map from 5kb resolution of Micro-C data, in which the coordination of ChIP peak was centered at the center point of each snippet. The snippets were then piled-up on the center of plot and normalized by the expected matrix.

i. **Motif analysis**

Sequences of loop anchors were extracted for CTCF cognate binding motifs scanning by MEME suit (http://meme-suite.org/). We also investigated the sequence enrichment for 20bp upstream and downstream of CTCF motif.

### List of Supplementary figures and tables

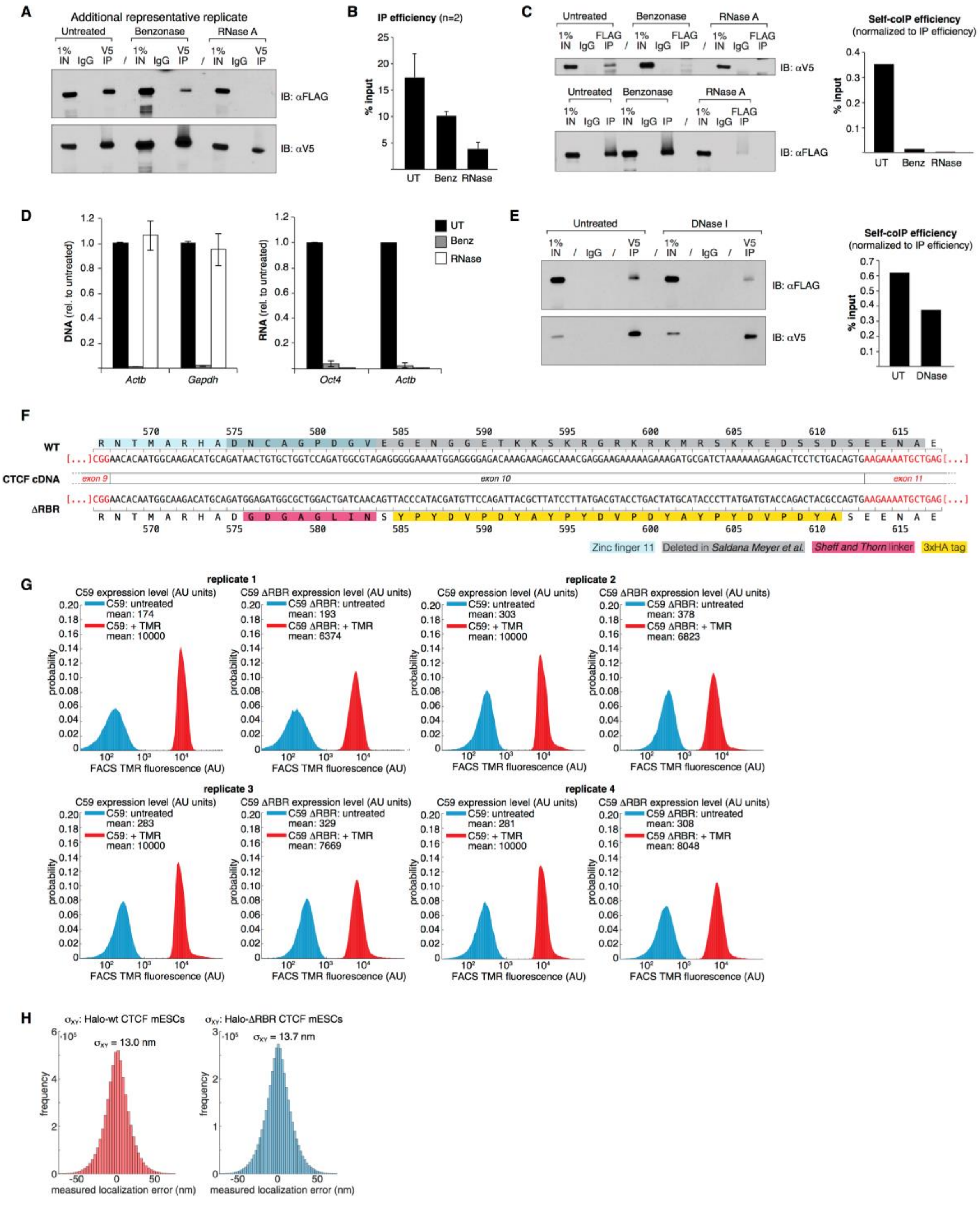


**Figure S1. Related to Figure 1 and 2.**

**Additional coIP experiments and controls**. (**A**) Representative coIP experiment (V5 IP) indicating RNA-dependent CTCF self-interaction (additional replicate). Top: V5 IP followed by FLAG immunoblotting measures self-coIP efficiency (90% of the IP sample loaded); bottom: V5 IP followed by V5 immunoblotting controls for IP efficiency (10% of IP sample loaded). (**B**) IP efficiency for the V5 coIP experiments of Figure 1C and S1A, used to calculate the self-coIP efficiency of Figure 1D. Error bars are SD, n=2. (**C**) Reciprocal coIP experiment (left) and quantification (right). Top left: FLAG IP followed by V5 immunoblotting measures self-coIP efficiency (45% of the IP sample loaded); bottom left: FLAG IP followed by FLAG immunoblotting controls for IP efficiency (45% of IP sample loaded). (**D**) Effective nucleic acid digestion by Benzonase (Benz) and RNase A during CTCF coIP experiments. DNA (left) and RNA (right) were extracted from coIP lysates and quantified by qPCR and RT-qPCR, respectively, using primers specific to *Actb* and *Gapdh* gene/mRNA. Error bars are SD, n=3. (**E**) coIP experiment with DNase I treatment (left) and quantification (right). Top left: V5 IP followed by FLAG immunoblotting measures self-coIP efficiency (80% of the IP sample loaded); bottom left: V5 IP followed by V5 immunoblotting controls for IP efficiency (20% of IP sample loaded). (**F**) Detailed view of CTCF mRNA exon 10 in wild type and ΔRBR-CTCF mESCs. Both the DNA and the protein sequences (one letter code) are provided. Relevant features are highlighted in colors. Amino acid numbers are based on the NCBI Reference Protein NP_851839.1.(**G**) Quantification of Halo-wt CTCF and C59 Halo-ΔRBR-CTCF expression levels in the mESC clones C59 and C59 ΔRBR, respectively. Cells were labeled with 500 nM Halo-TMR for 30 min and their background-subtracted fluorescence measured using on a LSR Fortessa Cytometer, exciting fluorescence with a 561 nm laser and collecting fluorescence through a 610/20 bandpass emission filter. All 4 biological replicates are shown. (**H**) Quantification of localization error/uncertainty in PALM experiments. Localization error (defined as the standard deviation) was measured from single-molecule localizations that appeared for at least 20 frames. We then calculated the mean X,Y coordinates and took the difference between the measured X,Y coordinates in a given frame from the overall mean to be the localization error.

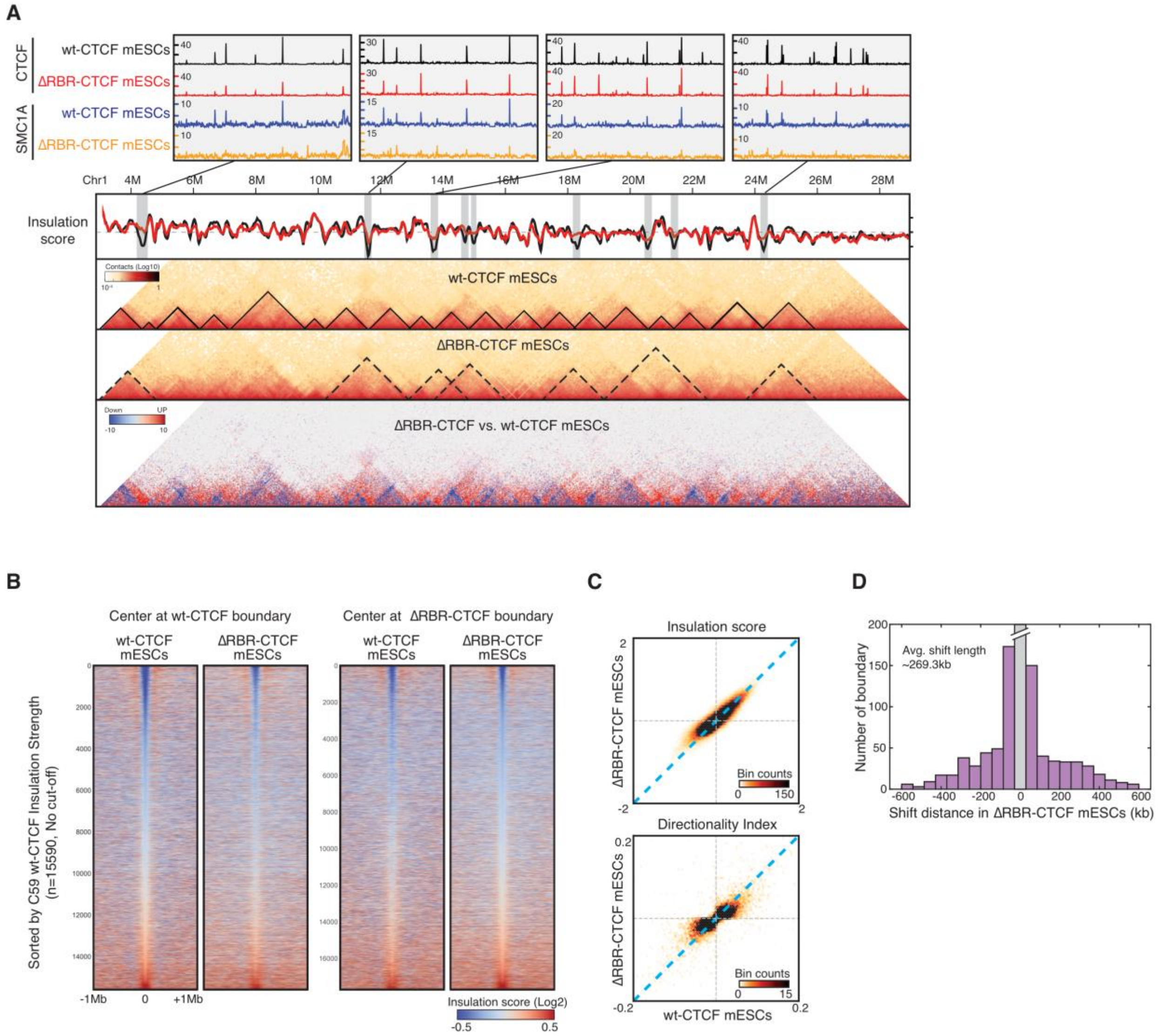


**Figure S2. Related to Figure 4.**

**TAD/Insulator analysis** (**A**) Additional example of TAD/Insulator disruption in ΔRBR-CTCF mESCs. Top panel shows the browser tracks of CTCF and Smc1a ChIP-Seq data zoomed-in on the regions where insulators were disrupted. Smc1a signal is largely reduced in ΔRBR-CTCF mESCs in all the cases shown here. Bottom panel shows a snapshot for insulation score, contact maps for wt-CTCF and ΔRBR-CTCF mESCs, and differential contact maps at Chr1:4M – 28M. Insulation scores were analyzed as described in the Methods section. A lower insulation score represents a strong insulator activity. ΔRBR-CTCF mESCs (red) lose some peaks comparing to wt-CTCF mESCs (black). Contact maps shown in the same region highlight larger TADs, which corresponds to the disruption of insulator in ΔRBR-CTCF mESCs. Gain of contacts between domains in ΔRBR-CTCF mESCs also supports this observation. (**B**) Heatmaps of insulation strength. Heatmap was centered at the called insulators (n=15590) in wt-CTCF or ΔRBR-CTCF mESCs flanking by ±1Mb and plotted by the rank of the insulation score from low to high (strong to weak insulators) without a cutoff value. The insulation strength is decreased in ΔRBR-CTCF mESCs while centering at the wt-CTCF insulators, with a greater degree in the group of the strongest insulators. However, there is no significant change in insulation strength between wt-CTCF and ΔRBR-CTCF mESCs while centering at ΔRBR-CTCF insulators. (**C**) Binned scatter plots of insulation score and directionality index. Scatter plots were binned to 250 bins and the quantity of each bin was shown as the color bar. Insulation score was analyzed as described in the Methods section and directionality index was analyzed as described in (Dixon et al., 2012). Two independent approaches confirmed that insulation strength is reduced in ΔRBR-CTCF mESCs. (**D**) Histogram of shift distance of ΔRBR-CTCF insulators. The cryptic insulators in ΔRBR-CTCF mESCs were taken to calculate one's distance to the closest insulator in wt-CTCF mESCs. The majority of cryptic insulators are shifted ~50 to 300kb from the original site (average length ~269.3kb). Insulators with shifted distance shorter than 40kb were precluded in the analysis.

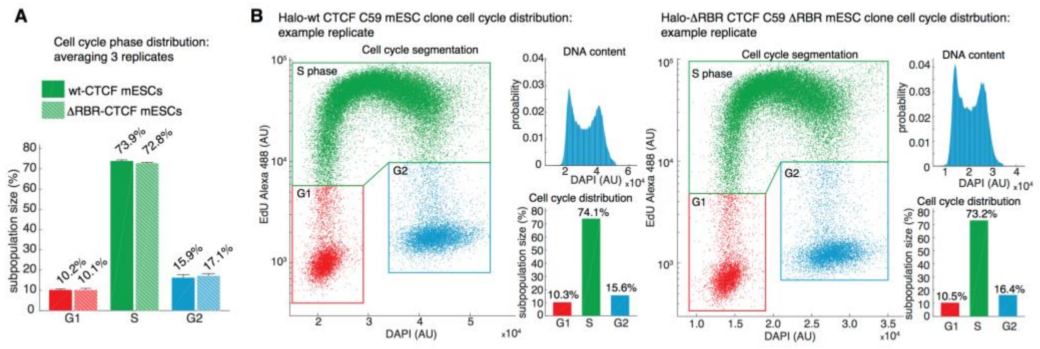


**Figure S3. Related to Figure 4.**
**No significant change to cell cycle distribution in ΔRBR-CTCF mESCs**. (**A**) Distribution of cell cycle phases in wt-CTCF and ΔRBR-CTCF mESCs (from 3 biological replicates). (**B**) Representative replicate (1 of 3). Cell cycle analysis was performed using the Click-iT EdU Alexa Fluor 488 Flow Cytometry Assay Kit (ThermoFisher Scientific Cat. # C10425) according to manufacturer's instructions. DNA content was inferred from DAPI staining and G1, S and G2 phases were gated as illustrated in red, green and blue in the plot.

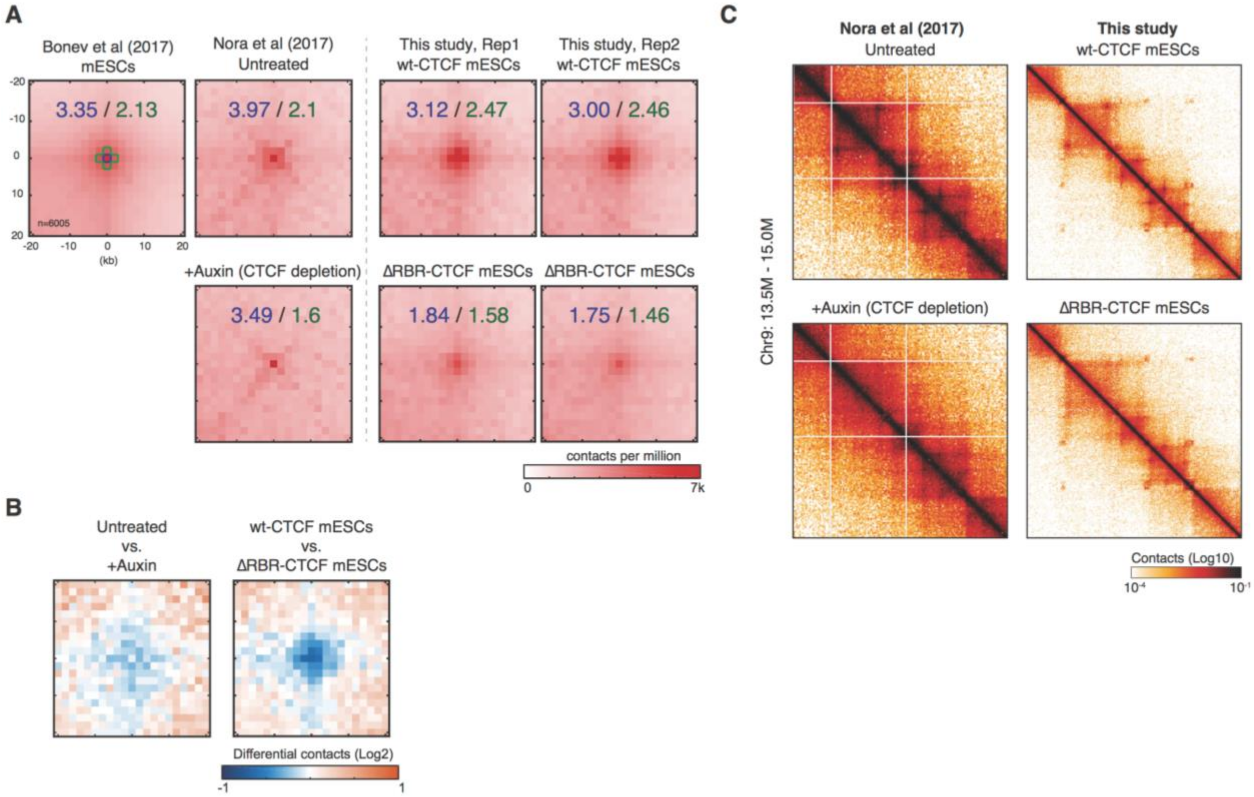


**Figure S4. Related to Figure 5.**

**Controls for loop analysis**. (**A**) Aggregate peak analysis for published Hi-C datasets and Micro-C. 6005 loops were called by using (Bonev et al., 2017) mESC dataset with FDR < 0.1. Datasets include mESCs in Bonev et al, wt-CTCF and AID-CTCF mESCs in (Nora et al., 2017), and two replicates of wt-CTCF and ΔRBR-CTCF mESCs generated by Micro-C. Loops were aggregated and plotted at the center of 20kb x 20kb window. The enrichments denoted on the plot were calculated by dividing one pixel at the center (blue) or five pixels around the center (green) by the average of bottom-left pixels. (**B**) Differential contact maps. Differential contact matrix was calculated by the log2 change between untreated vs. +auxin in (A) or wt-CTCF vs. ΔRBR-CTCF mESCs in (**A**). (**C**) Snapshots of the same region shown in Nora et al. Loops are largely disrupted in CTCF-depleted mESCs (left panel) and in ΔRBR-CTCF mESCs (right panel). This further confirms our genome-wide analysis that both CTCF-depletion and ΔRBR-CTCF affect loop formation/stability, although high-resolution Micro-C data may not be directly comparable to the standard Hi-C data.

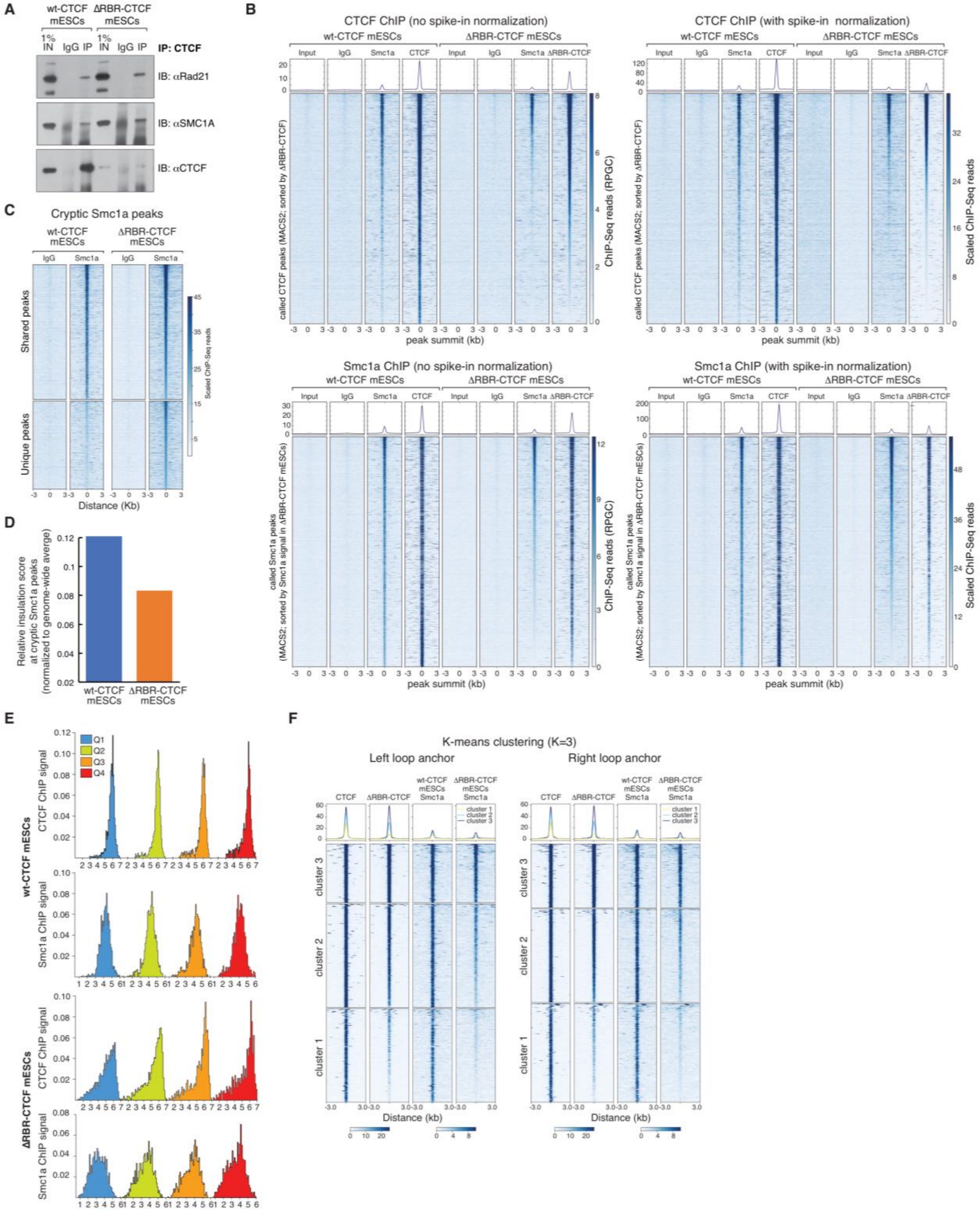

**Figure S5. Related to Figure 6.**

**Additional coIP experiment and ChIP-Seq / Micro-C analyses**. (**A**) Additional coIP experiment to the one shown in Figure 6A, demonstrating that CTCF interaction with cohesin is preserved upon deletion of the RBR. CTCF antibodies can pull down Rad21 and Smc1a cohesin subunits in both wt- and ΔRBR-CTCF mESCs. (**B**) Heatmaps of CTCF and Smc1a ChIP-Seq signal around wt-CTCF (top 2 panels) and Smc1a in wt-CTCF mESCs (bottom 2 panels) peaks as called by MACS2. We show results after normalization by sequencing depth (left panels; deepTools RPGC: reads per genomic content) or after normalization by the spike-in yeast DNA (right panels; scaled number of reads; see STAR Methods on how we computed the scale factor). Heatmaps are sorted by the peak intensity of ΔRBR-CTCF (top) and Smc1a in ΔRBR-CTCF mESCs (bottom). Above the heatmaps are summary plots with mean signal intensities. (**C**)

Heatmaps of IgG and Smc1a ChIP-Seq signal in wt-CTCF (left) and ∆RBR-CTCF (right) mESCs around Smc1a peaks called by MACS2 in ∆RBR-CTCF mESCs as shared or unique as described in Figure 6C. Read numbers are normalized by the spike-in yeast DNA (scaled number of reads; see STAR Methods on how we computed the scale factor). Heatmaps are sorted by ∆RBR-CTCF mESCs Smc1a signal intensity, and show that the peaks called as unique by MACS2 in ∆RBR-CTCF mESCs show very little, if any, enrichment of Smc1a in wt-CTCF mESCs. (**D**) Genome-wide normalized insulation score at the cryptic Smc1a peaks generated in ∆RBR-CTCF mESCs. Boundaries with a cryptic Smc1a peak within ± 40 kb were selected for insulation score analysis. Approximately 3.67% of the cryptic Smc1a peaks (646 out of 17,594) satisfied this criterion in ∆RBR-CTCF mESCs. Insulation scores for the individual boundaries were extracted and averaged, and further normalized to the genome-wide insulation score at the boundary (wt-CTCF=-0.417 and ∆RBR-CTCF=-0.336 (log2)). The normalized insulation score represents the relative enrichment of insulation strength, which presumably offsets the systemic shift in ∆RBR-CTCF mESCs (also see Figure S2B-C). A lower value of insulation score represents a higher insulation strength. (**E**) Histograms of ChIP signal distribution for Q1-Q4 loop anchors (also see CDF in Figure 6H). Loop anchors were identified as described in the Method section. ChIP signal enrichments were quantified in a 500 bp window centered at the anchors and plotted as a function of probability distribution in 50 bins. (**F**) Heatmaps of k-means clustering for the Q1 loop anchors (also see K-S probability density curve in Figure 6I). ChIP signals at ± 3 kb of the left and right loop anchors were clustered by k-means analysis with k=3 and sorted by the sum of regions in CTCF and Smc1a data in ∆RBR-CTCF mESCs. We also performed k-mean clustering analysis with k=2 and k=4 (not shown). Consistent to our conclusion, both results indicate two major subclasses of RBR-dependent structures with complete/partial loss (clusters 1 and 2, type 1 loop in Figure 5D) and no loss (cluster 3, type 2 loop in Figure 5D) of CTCF and/or cohesin in ∆RBR-CTCF mESCs.

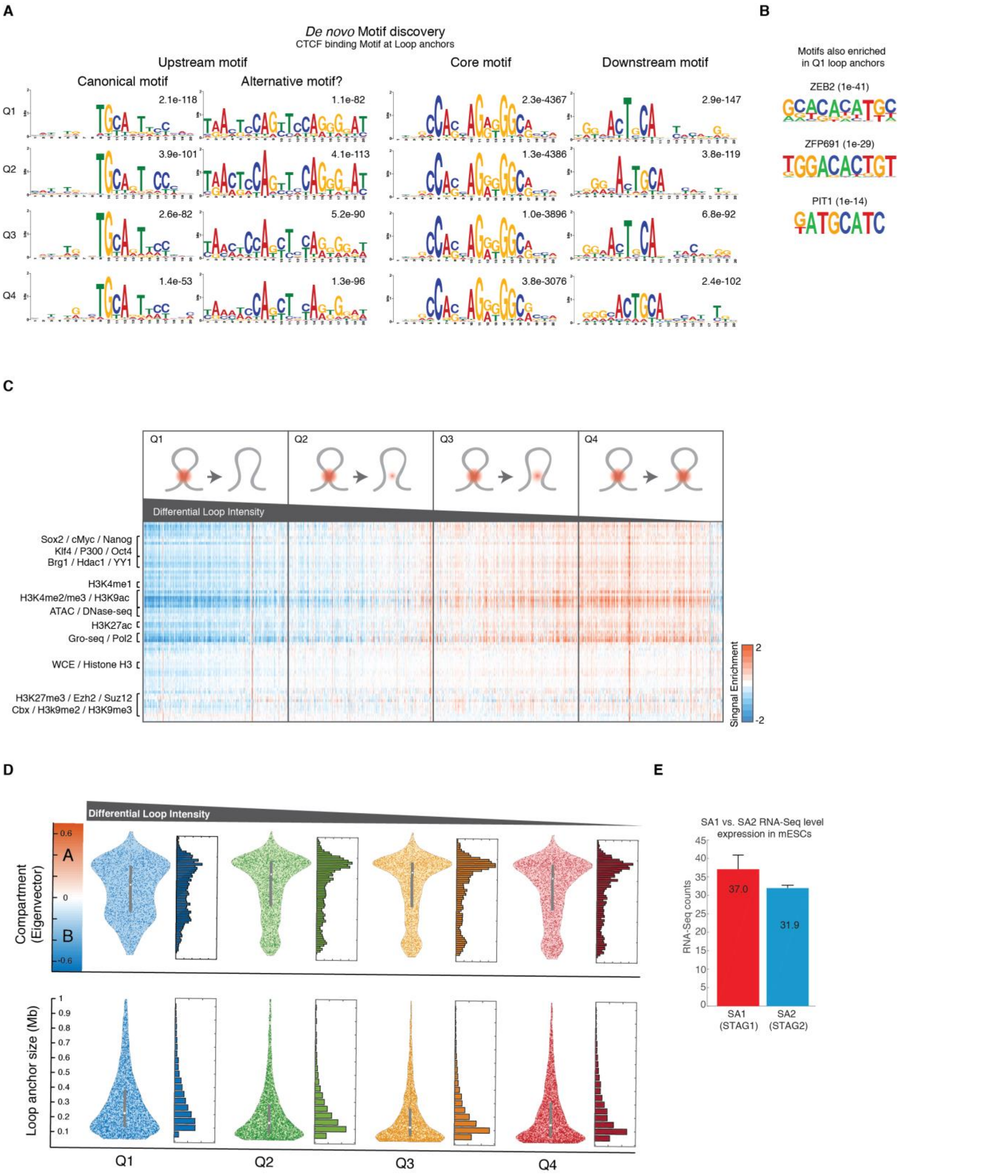


**Figure S6. Related to Figure 6.**

**DNA determinants of RBR-dependent and RBR-independent loops and potential factors in regulating loop formation/stability.** (**A**) *De novo* motif discovery at loop anchors. Loop anchors were identified as described in the Method section. Motif analysis was performed to identify potential binding sequences at the core loop anchors and ± 20 bp upstream and downstream of them. Motif searching was set as zero or one occurrence per sequence (zoops) mode by using MEME algorithm. Note that an alternative upstream motif was strongly enriched in addition to the CTCF canonical upstream motif, with an even greater significance (lower E-value) in Q2-Q4. The CTCF core motif was strongly and almost equally enriched at the anchors of chromatin loops Q1 through Q4 (Q1 > Q2 > Q3 > Q4), while the canonical CTCF upstream motif was particularly abundant at Q1 loops anchors. This is notable, since the upstream motif is the one recognized by CTCF Zn fingers 9-11 (Nakahashi et al., 2013), which are adjacent to the RBR domain. On the contrary, Q1 loop anchors, the least affected by RBR deletion, scored less significant for the canonical upstream motif, and were instead enriched in the alternative upstream motif. (**B**) Additional motifs discovered at Q1 loop anchors. Differential motif analysis was performed to identify potential factors involved in loop formation in addition to CTCF. Motifs were searched within a 5-kb window at loop anchors in Q1, and in Q4 as a control. Many prior uncharacterized zinc-finger proteins were shown with a high enrichment at loop anchors. These candidates can be subjects of future investigation. (**C**) Signal enrichments of 70 published genome-wide datasets at loop anchors were quantified as rlog variances by DEseq2. The heatmap was sorted by the changes in loop intensity between wt-CTCF and ΔRBR-CTCF mESCs. As a control, whole cell extract (WCE) and histone H3 has no significant change from Q1 to Q4. Most transcription-related factors (e.g. pluripotent factors, Brg1, YY1, etc.), active chromatin marks (e.g. H3K4me3, H3K9ac, etc.), enhancer marks (e.g. H3K4me1, H3K27ac, P300, etc.), and chromatin accessibility (e.g. ATAC-seq, DNase-seq) are strongly depleted in the Q1 loop anchors but enriched in the Q4 loop anchors. Constitutive heterochromatin marks (e.g. H3K9me2/3) and facultative heterochromatin marks (e.g. H3K27me3) have no preferential enrichment in any of the loop quartiles. (**D**) Distributions of loop anchors. Eigenvalues were ranked at the y-axis in top panel, in which positive $EV_1$ represents compartment A (active chromatin) and negative EV2 represents compartment B (inactive chromatin). More Q1 loops were found in inactive chromatin or at the border of two compartments but Q2 to Q4 loops were largely in active chromatin. Loop size distribution is similar across the four loop quartiles, ranging from 50 to 500 kb, although larger loops (> 500 kb) are enriched in Q1. This is consistent to a previous report that the average size of TADs/loops is larger in inactive chromatin. (**E**) SA1 and SA2 expression levels in mESCs. Plots show RNA-Seq levels (RPKM) from four replicates (GSE64040) from (Cattoglio et al., 2015).

**Table S1.**

**List of primer sequences.** This table contains a list of primers used in this study as well as a brief description of the experiment in which they were used.

| Name/description | Sequence (5'-3') | Experiment |
|---|---|---|
| Actb promoter forward | CATGGTGTCCGTTCTGAGTGATC | DNA quantification |
| Actb promoter reverse | ACAGCTTCTTTGCAGCTCCTTCG | DNA quantification |
| Gapdh transcription start site forward | TATCAGTTCGGAGCCCACAC | DNA quantification |
| Oct4 mRNA forward | CCAGGCAGGAGCACGAGTGG | DNA quantification |
| Oct4 mRNA reverse | CCTGGGACTCCTCGGGAGTTG | RNA quantification |
| Actb mRNA forward | GATCTGGCACCACACCTTCT | RNA quantification |
| Actb mRNA reverse | GGGGTGTTGAAGGTCTCAAA | RNA quantification |
| PCR primer 1 | AATGATACGGCGACCACCGAGATCTACACTCTTTCCCTACACGA | ChIP-Seq library prep |
| PCR primer 2 | CAAGCAGAAGACGGCATACGAGAT | ChIP-Seq library prep |
| C59D2 repair vector left homology arm forward | ATAGGGCGAATTGGGTACGGCATTTCCTTGCGTGGATTG | C59D2 Cas9 genome editing |
| C59D2 repair vector left homology arm reverse | GTTGATCAGTCCAGCGCCATCTCCATCTGCATGTCTTGCCATTGTG | C59D2 Cas9 genome editing |
| C59D2 repair vector right homology arm forward | GATGTACCAGACTACGCCAGTGGTAAGTGGGTCATTGTTG | C59D2 Cas9 genome editing |
| C59D2 repair vector right homology arm reverse | GGAACAAAAGCTGGAGCTAACCCGTGCAGATGGCTAAC | C59D2 Cas9 genome editing |
| C59D2 5' gRNA | CTGGTCCAGATGGCGTAGAG | C59D2 Cas9 genome editing; transfected alone or with the C59D2 3' gRNA |
| C59D2 3' gRNA | GACCCACTTACCACTGTCAG | C59D2 Cas9 genome editing; transfected with the C59D2 5' gRNA |
| CTCF intron 9 forward | AGAAATAAGATGACTTGAGTAGGCTG | C59D2 genotyping |
| CTCF intron 10 reverse | GTGAGTAGCACCTGGCTTTAG | C59D2 genotyping |
| Internal 3xHA | GGCGTAGTCTGGTACATCATAAGG | C59D2 genotyping |
| C62 gRNA#1 | ACTTACCAGAGACGCCGGGA | C62 Cas9 genome editing, V5-SNAPf tagging |
| C62 gRNA#2 | GGCCCCCTTCCCGGCGTCTC | C62 Cas9 genome editing, V5-SNAPf |

| | | |
|---|---|---|
| | | tagging |
| C62 gRNA#3 | AAAGACTTACCAGAGCGGGA | C62 Cas9 genome editing, 3xFLAG-Halo tagging |
| C62 gRNA#4 | TCCTGGCCCCCTTCCCGCTC | C62 Cas9 genome editing, 3xFLAG-Halo tagging |
| CTCF intron 2 forward | AGCACTGGTAGTTCTTTGTGGT | C62 genotyping |
| CTCF exon 3 reverse | GTTGGCTTCGGAGGCTTCATA | C62 genotyping |
| Internal SNAP | CTGTTCGCACCCAGACAGTT | C62 genotyping |
| Internal Halo | GTCGCGCTGGTCGAAGAATA | C62 genotyping |
